# Improvement in the enzymatic efficiency of a zinc metalloprotease through random mutagenesis approach: Studies on the enzymatic and structural properties of the mutant

**DOI:** 10.1101/2025.01.11.632526

**Authors:** Nitin Srivastava, Sunil Kumar Khare

## Abstract

Microbial proteases are significant enzymes acquiring 60% of the enzyme market. The heterologous expression provides higher enzymatic yield, yet such proteases possess less stability and low catalytic efficiency. A directed evolution strategy could improve protease’s properties to meet industrial demands. This study attempted the directed evolution of a recombinant metalloprotease (*rsep*) gene by a single round of error-prone PCR, wherein four hundred PCR were performed for random mutant gene library preparation. Eighteen amplicons (1230 bp) originated from modified PCR conditions were selected, cloned into pET22b+ plasmid, and overexpressed in *Escherichia coli* BL21(DE3). The recombinants showing soluble expression were screened, and one of the mutants (*rsep*A1) showed an 8.04-fold enhancement in the relative protease activity. It was purified to homogeneity with 92kDa molecular size. Its enzymatic properties like pH, temperature optima, and stability were comparable to *rsep*(WT). Furthermore, it exhibited improved enzymatic efficiency (∼ 4.2-fold towards casein) and better substrate affinity. The structural elucidation studies through bioinformatic tools, sequencing data, and comparative biophysical studies (Fluorescence-CD spectra-DSC) of the wild and mutant *rsep* showed increased histidine in the mutant. This probably resulted in enhanced substrate affinity and catalytic efficiency of *rsep*A1, which were well correlated with the ICP-MS analysis.

**Graphical Abstract:** 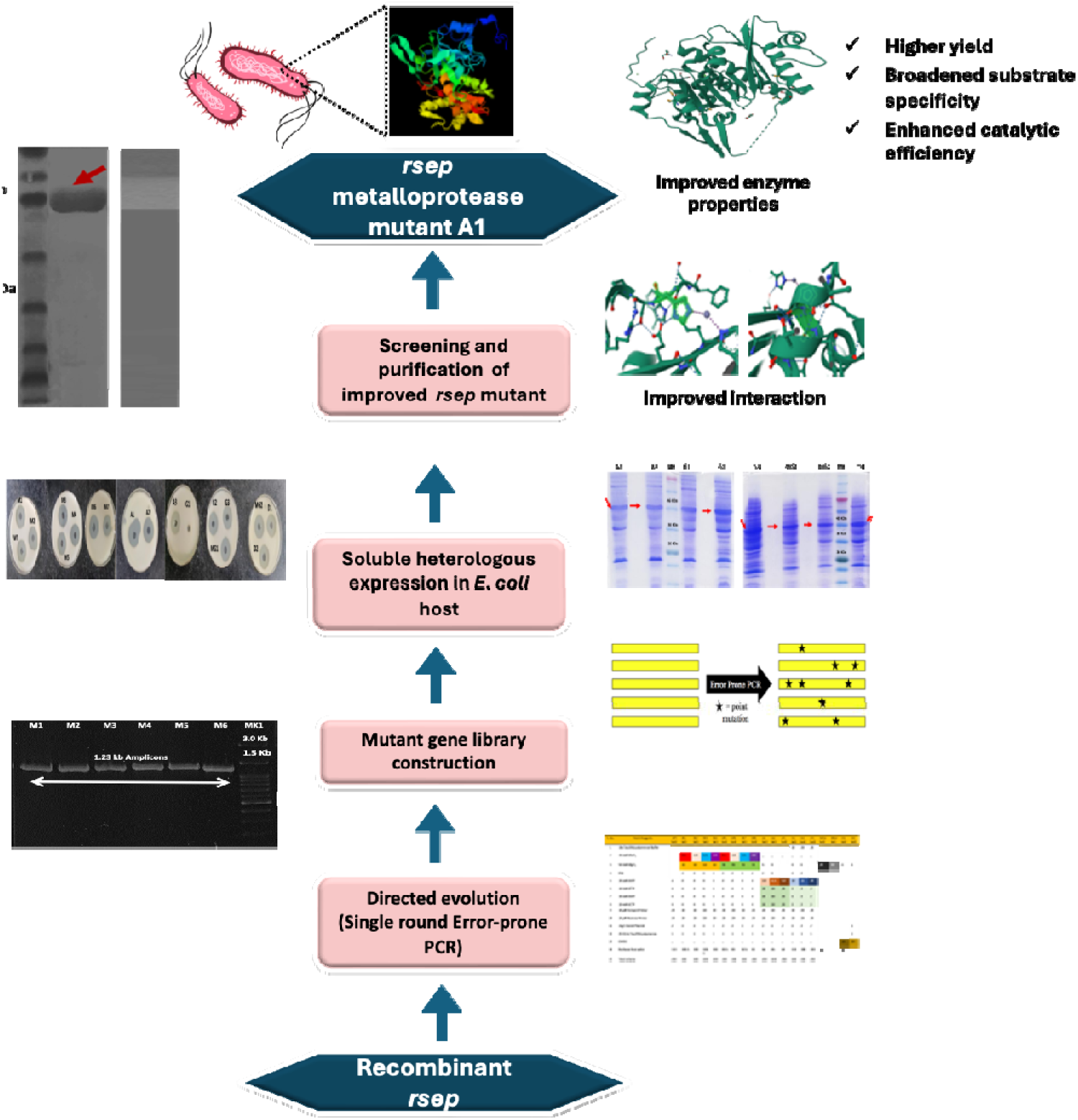

## 1. Introduction

Proteases are one of the major commercial enzymes. These possess versatile attributes, rendering them useful in various industries: bakery, brewing, feed, laundry, and detergent [1,2]. These also find applicability in processing silk, soy, and milk and are utilized for waste treatment and peptide synthesis. Such utilities allow them to acquire 60% of the total enzyme market [3,4]. Microbial proteases are more versatile and can be sufficiently produced in less time; they outrank plant and animal-derived proteases. Hence, these can satisfy the increasing demands of the various industrial sectors [5].

Most such proteases are produced through recombinant DNA strategies, which provide a relatively higher enzymatic yield, yet such proteases often have less stability and low catalytic efficiency. This limits their successful industrial implementations [6]. To achieve higher catalytic efficiency, better yields, and longer stability in a single protease, synthetic biology approaches to gene manipulation could be exploited. These techniques could help evolve and overexpress proteases with required attributes in the heterologous *E*. *coli* hosts [7,8].

The directed evolution approach could effectively tailor proteases of particular interest by employing random point mutations in their gene sequences. Directed evolution has been variously attempted to impart mutations in gene sequences of different enzymes of commercial interest. It could be implemented through chemical, liposome-mediated, microdroplets-based, DNA shuffling, site-directed, and Error-prone PCR-based random mutagenesis [7–10]. Some such attempted studies are designing and directed evolution of a commercial streptavidin through chemical methods [11], the directed evolution of alpha hemolysis by liposome display method [12], the cell-free directed evolution of a protease in microdroplets [13], enhancement of the thermostability of subtilisin E from *Thermus aquaticus* by the introduction of disulfide bond through site-directed mutagenesis approach [14], the directed evolution performed for enzyme epoxide hydrolase from *Agrobacterium radiobacter* through combinatorial approaches like error-prone (EP) PCR and DNA shuffling [15], the improvement of solvent tolerance property of keratinase from *Bacillus licheniformis* by directed evolution through error-prone PCR methodology [16], and the directed evolution of prolyl endopeptidase from *Flavobacterium meningosepticum* [17].

The error-prone polymerase chain reaction (EP-PCR)-based random mutagenesis strategy exploits the infidelity of Taq DNA polymerases to generate mutant gene libraries with various point mutations. It is a less complicated, cost-effective, less tedious, and equally efficient approach to developing improved enzymes with unknown structural details, better suited for industrial applications [7,8,18,19]. This process does not need as complex expertise as phage display, site-directed mutagenesis, and high-end computational approaches. Yet, it could lead to a product with suitably enhanced properties [7,19].

To our knowledge, the current study is the first report involving the improvement of a recombinant metalloprotease through a single round of EP-PCR. However, most similar studies focussed on improving the stability of the enzyme towards temperature and organic solvents for better functionality. This study used a previously cloned *rsep* metalloprotease gene cloned in pET22b(+) and attempted heterologous expression in the *E*. *coli* BL21(DE3) host [20]. The recombinant metalloprotease’s directed evolution, characterization, and structure elucidation were undertaken. A single round of EP-PCR-based mutagenesis approach has been employed to develop an improved mutant metalloprotease with enhanced enzymatic potential. Four hundred PCR reactions were performed for mutant gene library preparation. 18 modified, and a single control was suitably amplified at the predicted gene size of 1230 bp. The mutant and wild-type genes were cloned into the pET22b+ vector for overexpression. All the recombinants showed considerable expression in the soluble fraction. A screened mutant metalloprotease (*rsep*A1) showed an approximately 8.5-fold and 4.2-fold increase in protease activity and catalytic efficiency, respectively. It was purified to homogeneity with a molecular size of 92 kDa. The structural elucidation studies through bioinformatic tools, sequencing data, and comparative biophysical studies of the wild and mutant metalloprotease showed the presence of an increased amount of histidine residues in the mutant metalloprotease besides the appearance of acidic residues like glutamic, aspartic acid, etc. This led to a better interaction between the substrates and the mutant enzyme, which enhanced its affinity and catalytic efficiency, which was ensured by an increase in the amount of Zn^2+^ ions in the purified recombinant *rsep* metalloprotease compared to wild-type *rsep* metalloprotease through ICP-MS analysis.

## 2. Materials and methods

### 2.1. Materials

All the proteolytic substrates, reagents, and solvents utilized in the current study were obtained from Sigma-Aldrich (USA). The media components, salts, Ampicillin, and Bovine serum albumin (BSA) used in the present study were purchased from Hi-Media Laboratories Pvt. Ltd. (Mumbai, India). The Isopropyl-ß-D-thiogalactoside (IPTG) was procured from Sigma-Aldrich (USA). A 4 Å molecular sieve was used to treat all the solvents before using them for the experiments. The chemical constituents of polymerase chain reaction (PCR), such as Tris-HCl, Magnesium Chloride (MgCl_2_), Manganese Chloride (MnCl_2_), Dimethyl Sulphoxide (DMSO), and Potassium Chloride (KCl), were ordered from Invitrogen (by Thermo Fischer Scientific). dNTPs, restriction enzymes (BamHI and XhoI), Taq DNA polymerase, Quick CIP (Alkaline Phosphatase Calf Intestinal), and T4 λ-DNA ligase were purchased from New England Biolabs (NEB, USA). The primers were procured from IDT. All other solvents used for experimental purposes were ordered from Sigma-Aldrich and were of analytical grade. All the chemicals and reagents used for electrophoresis experiments were of molecular biology grade.

### 2.2. Bacterial strains, plasmids, and culture media

We previously cloned a metalloprotease gene (*rsep*-WP_195864791) from the genome of an *Exiguobacterium* sp. TBG-PICH-001 isolate into pET-22b(+) vector [21]. The cloned plasmid (pET22b(+)-*rsep*) was stored at -20°C. This study used the cloned plasmid as the template DNA for generating the mutant protease gene library. The *E*. *coli* strains DH5-α and BL21(DE3) (Novagen-Sigma Aldrich, India) were hosts for the recombinant plasmid’s heterologous expression. The *E*. *coli* strains were maintained on Luria Bertini (LB) agar media plates at 4°C, and their 50% glycerol stocks were stored at -80°C. The culture of *E*. *coli* was performed in LB media broth with 100 µg/ml ampicillin. The pET22b(+) plasmid vector was procured from Promega, USA. The LB media (containing 10 g/l tryptone, 5 g/l yeast extract, 10 g/l NaCl, 0.1 g/l CaCl_2_, 0.5 g/l ZnCl_2,_ 0.2 g/l KH_2_PO_4_) (adjusted to pH 7.0) was used for heterologous expression.

### 2.3. PCR primers

The forward (5’-CGGGATCCCGATGACGACCTTTATTTCAATCG-3’ [Tm,55°C]) and reverse (5’-CCCTCGAGGGAAAGAATTTTTGAATGTCGTTCC-3’ [Tm,55°C] primers were used for *rsep* gene amplification and library construction.

### 2.4. Library construction by error-prone PCR

The error-prone PCR methodology was used for random mutagenesis of *rsep,* followed by plasmid library construction [17]. The cloned plasmid (pET22b(+)-*rsep*) was used as the template. The constituents of the basic PCR (Table 1) were modified in various error-prone combinations (400 in number) to amplify *rsep.* The PCR was carried out using an MJ Mini Personal Thermal Cycler (Bio-Rad) with a program of 94°C for 1 min, 94°C for 30 s, 50°C for 30 s, and 72°C for 4 min (25 cycles), and finally, 72°C for 7 min. The resultant amplicons were screened based on the molecular size of the *rsep* gene (1230 bp) analyzed by horizontal gel electrophoresis using 1.5 % agarose. The agarose gels were visualized under ultraviolet light to observe the DNA bands analyzed in the gel documentation system (Bio-Rad, USA). The selected amplicons were digested with Dpn I at 37 °C for 1 h and gel-extracted using QIAquick Gel Extraction Kit (Qiagen, Germany) after electrophoresis through 1.5 % agarose gels.

**Table 1.**
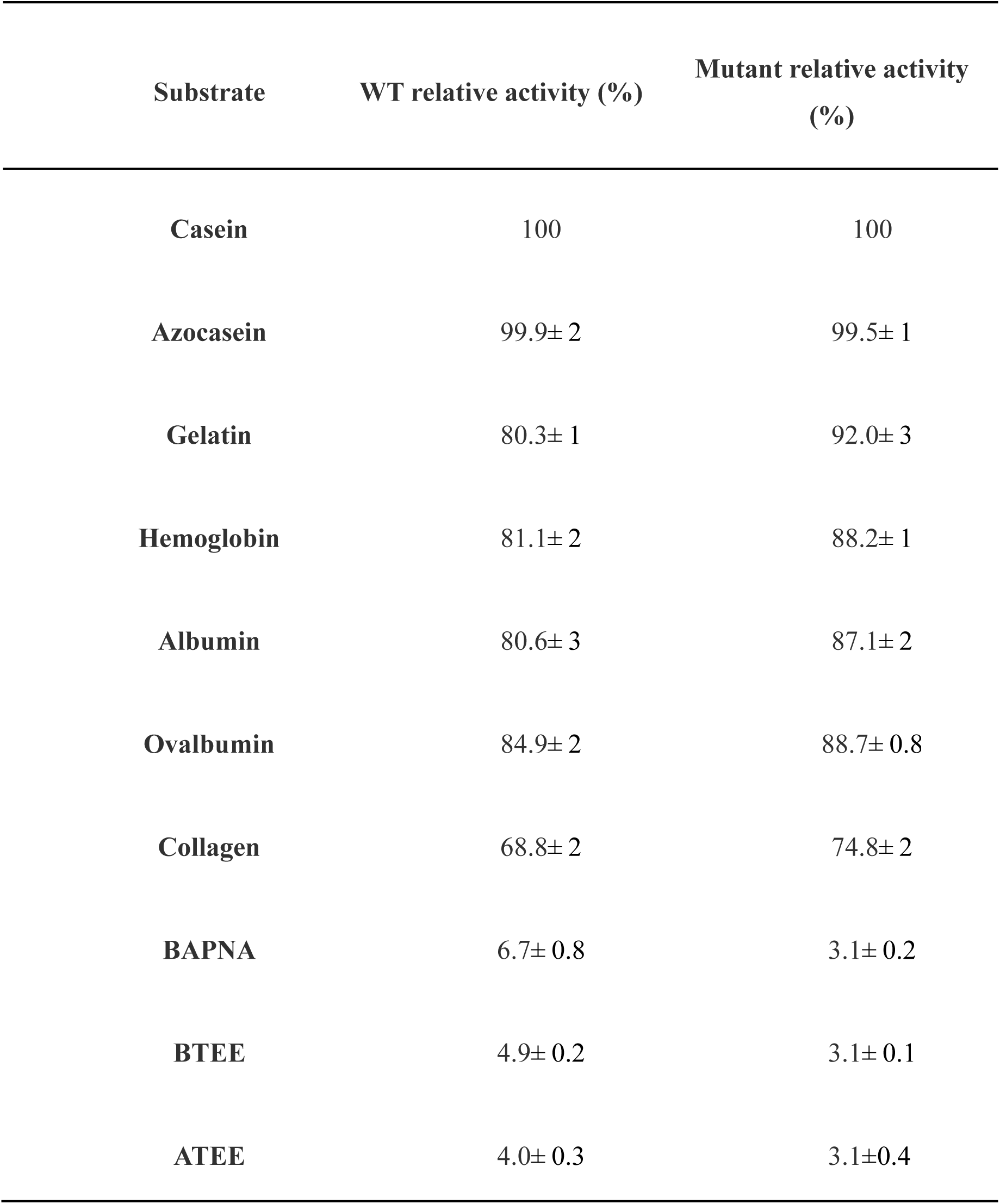
Substrate specificity studies of purified *rsep* mutant metalloprotease

### 2.5. Cloning and construction of plasmid library of mutant rsep genes

The gel-extracted *rsep* amplicons and pET22b+ plasmid vector (Promega, USA) were double digested with Bam HI and Xho I restriction endonucleases. The digested vector and amplicons were ligated using T4 DNA ligase (1U) for 16 h at 18 °C. The ligation mixtures were chemically (CaCl_2_ method) transformed in *E*. *coli* strain DH5-α competent cells. The transformed cells were spread-plated on the LB media agar plates containing ampicillin (100 μg/ml) as a resistant marker and incubated for 12 h at 37 °C [22]. The recombinant colonies on the overnight LB media plates were picked and transferred into 5 ml of LB media broth containing ampicillin (100 µg ml^-1^). These were incubated at 37°C and 180 rpm. The recombinant plasmids containing wild-type and mutant *rsep* were isolated using the alkaline lysis protocol [23]. These plasmid DNA samples (∼200 ng) were digested in a 20 μl reaction mixture with Bam HI and Xho I for 1 h at 37°C. The digested samples and standard size markers (1 Kb and 100 bp size) were resolved on 1 % agarose gel to analyze the restriction patterns and confirm the precise cloning. All the confirmed clones (plasmids) with wild-type and mutant *rsep* were labelled and stored at -20°C as a plasmid library of *rsep* mutants.

### 2.6. Overexpression analysis of rsep mutants

The recombinant wild-type and mutant (cloned plasmids) of *rsep* gene products were transformed into the host *E*. *coli* BL21(DE3) competent cells and spread-plated on LB media agar plates. The plates were incubated at 37°C for 12 h. A single colony for all the recombinants was separately inoculated in a 5 ml Luria Bertini (LB) media and incubated overnight at 150 rpm. A 1% v/v inoculum from the primary cultures of host *E. coli* containing recombinant plasmids was inoculated into separate and fresh 10 ml LB media (containing 10 g/l tryptone, 5 g/l yeast extract, 10 g/l NaCl, 0.1 g/l CaCl_2_, 0.5 g/l ZnCl_2,_ 0.2 g/l KH_2_PO_4_) broth (adjusted to pH 7.0). After the cultures reached an optical density (OD_600_) of 0.60, the cells were induced with 0.6 mM isopropyl-ß-D-thiogalactoside (IPTG) and incubated at 37°C, 180 rpm of constant shaking for 4 h. The cells were lysed using the lysis buffer (50 mM Tris-HCl, pH 8.0) and sonication (Amplitude 32 W; On pulse, 12s; Off pulse, 8s; total time, 40 mins). After the cell lysis, the cell lysate and sonicated protein supernatant samples were run on a 12% SDS-PAGE to check the protein expression profile for all the recombinant proteases. A 0.1% Coomassie Brilliant Blue R250 in water: methanol: glacial acetic acid solution (4.5:4.5:1) was used for staining the polyacrylamide gels. The stained gels were de-stained using water: methanol: glacial acetic acid solution (4.5:4.5:1). The molecular weight was determined by comparing the protein profile with a pre-stained protein molecular weight standard marker [22].

### 2.7. Screening of mutant rsep proteases

The cloned mutants and the wild-type *rsep* were expressed in the *E. coli* BL21(DE3), and their cells were harvested for the comparative estimation of protease activity through qualitative and quantitative analysis. In the qualitative analysis, the soluble protein fraction was utilized for comparative estimation of the hydrolytic zone in the mutants. In contrast, the quantitative analysis estimated recombinant wild-type and mutants’ comparative protease activity (calculated in standard units, i.e., IU).

#### 2.7.1. Nutrient broth-agar-well diffusion assay

The nutrient broth, agar-agar, and gelatin (2% w/v) were used, and media agar plates were constituted. The media plates were bored to make plugs, and the soluble protein fractions (containing the expressed recombinant proteases) were poured and sealed with agar to form agar plugs in the plates. These plates were incubated at 37 °C overnight, and different samples were analyzed for the hydrolytic zones, which were compared with the recombinant wild-type to select the mutant with enhanced protease activity.

#### 2.7.2. Protease activity assay

Protease activity was measured using the method of Shimogaki *et al*. [24]. Briefly, 0.5 mL of the enzyme was added to 3mL casein 0.6 % w/v in 50 mM Tris-HCl, pH 9.0, and the reaction mixture was incubated at 50 °C for 20 min. The reaction was stopped by adding 3.2 mL of TCA mixture (0.1 M trichloroacetic acid, 0.2 M sodium acetate, and 0.3 M acetic acid), followed by keeping the reaction mixture at room temperature for 30 min. The precipitates were removed by filtration using Whatman-1 filter paper, and the absorbance of filtrate was measured at 280 nm. One unit of protease activity was defined as the enzyme amount required to liberate one micromole of tyrosine per minute under assay conditions. The mutants and the recombinant wild-type *rsep* were compared for the relative protease as per the standard protease assay [24].

### 2.8. Purification of the recombinant mutant proteases

The selected mutant protease A1 (with enhanced protease activity) was purified through the Ni-NTA affinity column (Qiagen, Germany) under non-denaturing conditions and analyzed on 12% SDS-PAGE according to the protocol described by Sambrook and Russell [22]. Briefly, the recombinant *E*. *coli* BL21(DE3) strain containing the cloned mutant *rsep* gene in the pET 22b(+) plasmids was inoculated in an LB media overnight at 150 rpm. A 1% v/v inoculum from the overnight grown culture of the mutant was introduced in a 200 mL LB broth (containing 10 g/l tryptone, 5 g/l yeast extract, 10 g/l NaCl, 0.1 g/l CaCl_2_, 0.5 g/l ZnCl_2,_ 0.2 g/l KH_2_PO_4_). The culture was grown at 37 °C at 180 rpm and induced with a 0.6 mM filter sterilized IPTG solution. The uninduced sample was removed from the cultures before induction and was separately grown for further periods. Both uninduced and induced cells were grown for another 4 h. The cells from the culture were harvested. These were centrifuged at 10,000 rpm for 10 minutes and resuspended in 15 mL lysis buffer (50 mM Tris-HCl, pH 8.0). They were disrupted by sonication (Amplitude 32 W; On pulse, 12s; Off pulse, 8s; total time, 40 mins), followed by centrifugation to remove the debris. The soluble expression in the supernatant was checked and used for further purification through His-tagged-based Ni-NTA affinity column chromatography. The His-tagged proteins were eluted with a gradient of imidazole concentration ranging from 100-500 mM. All the eluted fractions were checked for purification on the 12 % SDS-PAGE gel. The purified fractions were pooled and stored at -20 °C for further studies.

### 2.9. Estimation of protein concentration, SDS-PAGE, and zymography

The protein concentration was determined by a dye-binding assay described by Bradford using BSA as a standard protein [25]. The protein solution (0.5 ml) was incubated with 4.5 ml of the dye reagent for 5 min, and the absorbance of the solution was recorded at 595 nm against buffer blank (0.1M Tris-HCl buffer, pH 8.0). The calibration curve was plotted using BSA as a standard protein.

SDS-PAGE of the purified protease was carried out using 12% polyacrylamide gel on a Mini-PROTEAN 3 system gel electrophoresis unit (BioRad Laboratories, USA). For zymography, the samples were electrophoresed and stained for protease activity as per the modified method of Garcia-Carreno *et al*. [26]. Briefly, the electrophoresed gels were immersed in 100 mL of 2% (w/v) gelatin solubilized in phosphate buffer (0.02 *M*, *p*H 7.5 containing 1%, w/v NaCl) and incubated for 1 h at 37 ^◦^C. The gels were then fixed, stained with 0.125% (w/v) Coomassie blue R-250 (in 50%, v/v ethanol and 10%, v/v acetic acid), and destained with a solution containing methanol, acetic acid, and water (30:10:60). The development of clear zones against a dark background indicated the presence of protease activity.

### 2.10. Comparative characterization of the developed rsep mutant protease with the wild-type recombinant rsep

The mutant *rsep* was compared for its functional and structural properties with the wild-type recombinant *rsep*. These properties included optimum pH, temperature, pH & temperature stability, substrate specificity, kinetic parameters, the effect of metal ions, surfactants, and protease inhibitors, and solvent stability. Further, the mutant and wild-type recombinant proteases were compared for the modifications in the nucleotide and the consequent amino-acid residues through the sequencing data. The recombinant cloned mutant and wild-type plasmid sequencing was outsourced from Barcode Biosciences (Bangalore, India). Structural analysis was performed on these proteases to correlate these changes through fluorescence and circular dichroism spectral studies.

#### 2.10.1. Enzymatic properties

The effect of pH on pure enzyme was studied by assaying it at different pH. The pH stability was studied by pre-incubating the enzymes in buffers of different pH values (0.1M acetate buffer (pH 4.0-6.0); 0.1M phosphate buffer (pH 6.0-8.0); 0.1M Tris-HCl buffer (pH 8.0-9.0), and 0.1M Glycine-NaOH buffer (pH 9-10) was used), at 37 ^◦^C for 24 h. The residual activities were determined under standard assay conditions [27].

The temperature optimum for the purified protease was determined by performing the assay at various temperatures using 0.6% (w/v) casein as the substrate. The assay conditions were the same as described in the previous section. The thermal stability was studied by incubating the enzymes at 20-80 ^◦^C. The aliquots were withdrawn at different intervals, and the residual activities were determined under standard assay conditions. The pH conditions for performing these experiments were pH optima of the enzyme.

The specificity of the proteases was determined towards various protein substrates (such as casein, azocasein, albumin, ovalbumin, collagen, keratin, etc.) esters, and synthetic peptides conjugated with *p*-nitroanilide (pNA) by following the standard assay procedures. The substrate for which the proteases showed the highest activity was considered to have a 100% relative protease activity.

The kinetic parameters Michaelis-Menten constant (K_m_) and the maximum rate (V_m_) of the enzyme activity related to the enzyme were calculated through double reciprocal Lineweaver Burk plot equation of protease activity towards casein as substrates at varying concentrations. The substrate concentrations ranged from 0.016 mM-1 mM (0.16, 0.024, 0.032, 0.04, 0.08, 012, up to 1 mM) of the substrates like casein, azocasein, and gelatin [27].

The effect of metal ions, surfactants, and protease inhibitors on the enzymes was checked. The enzymes were incubated with different chemicals at varying concentrations at 37 ^◦^C for 1 h for metal ions like Zn^2+^, Ca^2+^, Mg^2^+, Pb^2+^, Ni^+^, Cu^2+^, Fe^2+^, K^+^, Hg^+^, and Mn^2+^, 30 min in case of surfactants (Tween 80, Triton X-100, Sodium dodecyl sulphate (SDS), and Urea) and inhibitors (β-Mercaptoethanol, Glutathione, Dithiothreitol (DTT), Iodoacetamide (IAA), p-Aminobezidine, PMSF, 1,10-Phenanthroline, Ethylenediaminetetraacetate (EDTA). The residual activities were measured as per the described standard assay procedure [28].

The organic solvent stability of the purified mutant protease was estimated. A known amount of purified mutant protease was dissolved in Tris-HCl buffer (0.1M; pH 8.5). A 3 mL of a known amount of each enzyme was separately mixed with 6 mL of various organic solvents (tetradecane, dodecane, decane, heptane, octane, isooctane, hexane, dichloromethane, and butanol). The mixture was sealed in parafilm and incubated at 200 rpm with constant shaking at 37 ° C for 48 h. The sample aliquots of equal volumes were collected from the mixture at 6, 12, 24, and 48 h. The organic solvent untreated sample remained the control and was incubated under similar conditions. As mentioned, the samples were checked for residual protease activity [28].

#### 2.10.2. Structural elucidation

The modifications in the nucleotide and amino acid sequences of the mutant *rsep* compared to the recombinant wild-type *rsep* were analysed using Expasy translate and the multiple sequence alignment tool (Clustal W online software). The point mutations were identified in the mutant protease sequences. Further, the sequence data of the mutant was correlated with its protein structure. A known amount of the purified protease was used to check the differences in the enzyme structure of the mutant protease as compared to the recombinant wild-type. The enzymes’ circular dichroism (CD) and fluorescence spectra were recorded for their secondary and tertiary structure analysis.

##### 2.10.2.1. Fluorescence spectroscopy

Fluorescence spectra were acquired on model Fluoromax-4 spectrofluorometer (Horiba-Jobin Yvon, Inc., NJ, USA) at room temperature using 1 cm^2^ path length quartz cuvettes. The excitation wavelength was kept at 295 nm. Emission spectra were recorded between 305 and 400 nm. Measurements were done in Tris-HCl buffer (0.1M; pH 8.5) at a protein concentration of 50 μg mL^−1^. Baselines were corrected with the corresponding buffers. The data recorded in triplicate were averaged and analyzed using Origin 8.0 software (USA) [29].

##### 2.10.2.2. Circular dichroism spectroscopy

The UV-CD spectra of the protease were recorded between 195 and 260 nm on a Chirascan Spectropolarimeter (Applied Photophysics, Leatherhead, Surrey, UK) at 20 °C using 1 mm quartz cuvette. An average of three independent scans was used. All measurements were performed in Tris-HCl buffer (0.1 M; pH 8.5) and at 0.5 mg mL^−1^ protein concentration. The contribution of respective buffers was subtracted from experimental spectra and further smoothed using a mild smoothing function [29].

### 2.11. Inductively coupled plasma mass spectrometry (ICP-MS) analysis

A 1 mg of the purified wild-type and mutant *rsep* metalloprotease were incubated in a 3% (v/v) HNO_3_ solution at 30 °C and 180 rpm for 48 h [30]. The debris was separated from the solution, and the enzyme-free solution was analyzed to estimate metal ions using inductively coupled plasma mass spectrometry (ICP-MS) [31].

## 3. Results and discussion

### 3.1. Development of novel rsep mutants through the directed evolution approach via. a single round error-prone (EP) PCR

The exploitation of the infidelity of the Taq DNA Polymerase could be an efficient technique to insert point mutations at a sufficient rate into a gene of interest in the presence of a modified chemical constitution of the PCR reaction. This could be instrumental in generating random mutant gene libraries in simple expression plasmids to attain improved enzymes (directed evolution), which can be utilized for industrial applications inexpensively [10,19,32]. In this context, the *rsep* mutant gene library was developed through a directed evolution approach via error-prone polymerase chain reaction (EP-PCR) using pET22b-*rsep* cloned plasmid DNA (containing wild recombinant *rsep* gene) as a parent template for setting up EP-PCR reactions. Only a single round of EP-PCR was employed for gene amplifications. The suitably designed oligonucleotide primers were used for the *in-vitro* synthesis of evolved *rsep* gene mutant through PCR amplifications that resulted in approximately1.23 Kb gene size amplification in 18 EP-PCR (M1-M8, MG1-MG2, D1-D2, A1-A3, G1-G3) conditions and a control reaction (C) condition **(Fig. 1)**. The gene amplification bands in the EP-PCR conditions were similar in size compared to the control (The primary PCR reaction with basic concentrations of the constituents (control) and the EP-PCR reactions with modified constituent concentrations are described in **Table S1** and the EP-PCR (modified) reactions for *in vitro* generation of evolved *rsep* mutant genes are described in **Table S2)**.

**Fig. 1.**
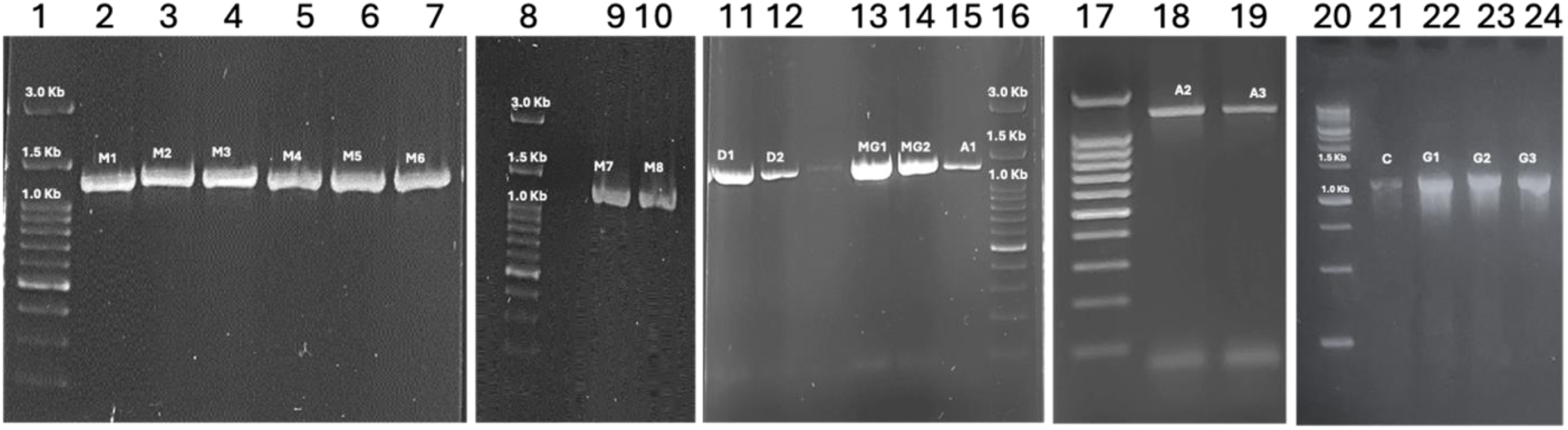
PCR-amplified ORF fragments of recombinant wild-type *rsep* metalloprotease and mutant *rsep* metalloprotease genes. Ethidium bromide-stained gels show the presence of 1.23 Kb fragments of *rsep* metalloprotease genes in basic PCR (Wild-type (WT) gene) and error-prone (EP)PCR conditions (Error-prone genes). Lane 1, 100 bp DNA ladder; Lanes 2,3,4,5,6,7,9,10, PCR amplified samples of Mutants M1, M2,M3,M4,M5,M6,M7,M8,respectively; Lane 8, 100 bp DNA ladder; Lanes 11,12, PCR amplified fragments of Mutants D1, D2; Lanes 13,14,15, PCR amplified samples of Mutants MG1,MG2,A1; Lane 16,17,100 bp DNA ladders; Lanes 18,19, PCR amplified samples of Mutants A2,A3; Lane 20, 100 bp DNA ladder; Lanes 21, Control; Lanes 22,23,24, PCR amplified samples of mutants G1,G2,G3. WT-Wild type *rsep* metalloprotease gene. M1, M2, M3,M4,M5,M6,M7,M8,A1,A2,A3,G1,G2,G3,MG1,MG2,D1,D2-(EP)PCR mutant genes. The mutant gene products were named after the modified conditions in the (EP)PCR M1:Mutant developed in 25 µM MnCl_2_, 5.0 mM MgCl_2_; M2:Mutant developed in 50 µM MnCl_2_, 5.0 mM MgCl_2_; M3:Mutant developed in 75 µM MnCl_2_, 5.0 mM MgCl_2_; M4:Mutant developed in 100 µM MnCl_2_,5.0 mM MgCl_2;_ M5:Mutant developed in 25 µM MnCl_2_,10 mM MgCl_2_; M6: Mutant developed in 50 µM MnCl_2_,10 mM MgCl_2_; M7:Mutant developed in 75 µM MnCl_2_, 10 mM MgCl_2_, M8:Mutant developed in 100 µM MnCl_2_,10 mM MgCl_2_; A1:Mutant developed in 0.15 mM dATP, 1.0 mM other dNTPs; A2:Mutant developed in 0.20 mM dATP, 1.0 mM other dNTPs; A3:Mutant developed in 0.25 mM dATP, 1.0 mM other dNTPs; G1:Mutant developed in 0.5 mM dGTP, 0.1 mM other dNTPs; G2:Mutant developed in 0.75 mM dGTP, 0.1 mM other dNTPs; G3:Mutant developed in 1.0 mM dGTP, 0.1 mM other dNTPs; MG1: Mutant developed in 5 mM MgCl_2_; MG2: Mutant developed in 10 mM MgCl_2_; D1:Mutant developed in 5 % v/v dimethylsuphoxide; D2:Mutant developed in 10 % v/v dimethylsulphoxide.

Various researchers have performed similar attempts at enzyme evolution for *in-vitro* generation by EP-PCR to derive random enzyme mutants with improved functionality. One such research deals with improving the thermostability of a prolyl endopeptidase gene from *Flavobacterium meningosepticum* through a directed evolution approach. They could produce the most thermostable mutant after three repeated EP-PCR cycles using an expression plasmid, pUK-FPEPb, as the master DNA template for PCR reactions [17]. Another similar research culminated in a successful attempt to generate an evolved epoxide hydrolase (EchA) gene from *Agrobacterium radiobacter* with higher enantioselectivity using a single round of EP-PCR reaction [15]. One of the studies investigated the PCR-based random mutagenesis using manganese and reduced dNTP concentration to check the effectivity and accuracy of the process to incorporate point mutations. The gene under investigation in this study was DNA mismatch repair protein (Msh2) [32]. A recent study similarly employed the EP-PCR for random mutagenesis to improve enzymes alcohol dehydrogenase and cyclohexanone monooxygenase. The improved enzymes were cloned in host *E*. *coli* and utilized as cell factories to produce ε-caprolactone [33].

The EP-PCR-based *rsep* mutants were used for restriction double digestion and ligated with the similarly digested pET22b+ plasmid to construct mutant *rsep* libraries.

### 3.2. Construction of mutant rsep plasmid libraries

All these 19 genes were cloned into the pET22b+ expression plasmid vector. The appearance of the colonies on the LB-agar media plates containing ampicillin (100 µg/ml) showed proper ligation and transformation of the genes and ligated plasmids, respectively. The single round of EP-PCR reaction and infidelity rate of Taq-DNA polymerase enzyme necessitated an intensive screening of bacterial colonies that may possess the precise ligation of mutant genes with point mutations; hence, many sets of mutant gene ligation were performed, and all possible bacterial colonies were checked for accurate amplification of the concerned gene’s molecular size. Out of the total colonies, 30-40 bacterial colonies, on average, showed precise amplifications in colony PCR. These colonies were used to isolate mutant plasmids for each mutant condition. All these plasmids were stored at -20 °C. Likewise, mutant gene libraries were created for all the mutant *rsep* generated by EP-PCR in pET 22b+ plasmid vectors. This resulted in approximately 30-40 cloned mutant plasmids for each mutant *rsep*. These were all subjected to restriction digestion using Bam HI and Xho I to confirm the precise cloning of the mutant *rsep*. A total of 15-20 cloned plasmids showed gene fallouts of correct molecular size (1230 bp), considering all the mutant *rsep* libraries **(Figure S1)**.

Many researchers carried out similar studies. In a study, mutant prolyl endopeptidase libraries were created in an expression plasmid, pUK-FPEPb transformed in host *E*. *coli* JM109 strain [17]. One such study used pBADmycHisA plasmids to develop libraries of enzyme Epoxide hydrolase. These plasmid libraries with different evolved gene mutants were transformed into host *E*. *coli* Top10 strains for further screening of enzyme improvement [15]. In a similar study, the researchers synthesised evolved circularised genes using a microdroplet-based approach introduced in pHAT plasmids to construct savinase (protease) gene libraries for their PCR-mediated directed evolution and screening for functional improvement [13]. In another research, disulfide mutants were generated through a site-directed mutagenesis approach for enhancement of the thermostability of Subtilisin E, where mutant genes library were created by transforming these in different plasmid vectors (pHTC61, pHTC98, pHTC61C98) to develop plasmid gene libraries with specifically mutated amino acid residues [14].

These plasmid libraries with appropriately cloned *rsep* gene mutants were selected to investigate the heterologous expression and protease activity further.

### 3.3. Heterologous expression of the rsep mutants in host E. coli and their comparative protease activity

The screened mutant *rsep* plasmids were transformed into the host *E*. *coli* BL21(DE3) cells. A significant number of colonies for all the generated *rsep* mutants were investigated for overexpression of the *rsep* mutants. The colonies were grown in modified LB media (production media). The harvested cells were lysed and checked for overexpression in soluble and inclusion body protein fractions. All the colonies demonstrated accurate expression of the *rsep* in the soluble protein fraction with a molecular size of 46 kDa by denaturing PAGE analysis **(Fig. 2a)**. The enzyme expression was also observed in the inclusion body fraction but much less than that of the soluble enzyme **(Fig. 2b)**. Hence, the soluble protein fraction (crude enzyme) was utilized to check their comparative proteolytic activity with the wild-recombinant *rsep*. A similar study demonstrated the over-expression of prolyl endopeptidase in the *E*. *coli* JM109 strain. The host cells containing expressed enzyme were lysed to separate the soluble enzyme fraction, which was investigated for further screening and selection of improved mutant enzymes [17]. Several other successful studies have been conducted on the heterologous expression of proteases from microbial genomes and environmental DNA samples. This includes protease genes from environmental DNA samples from coastal Gujarat [34], *Haloakalobacillus lehensis* [35], *Nocardiopsis* sp. [36], *Pseudomonas aeruginosa* K [37], and *Bacillus lehensis* [38].

**Fig. 2.**
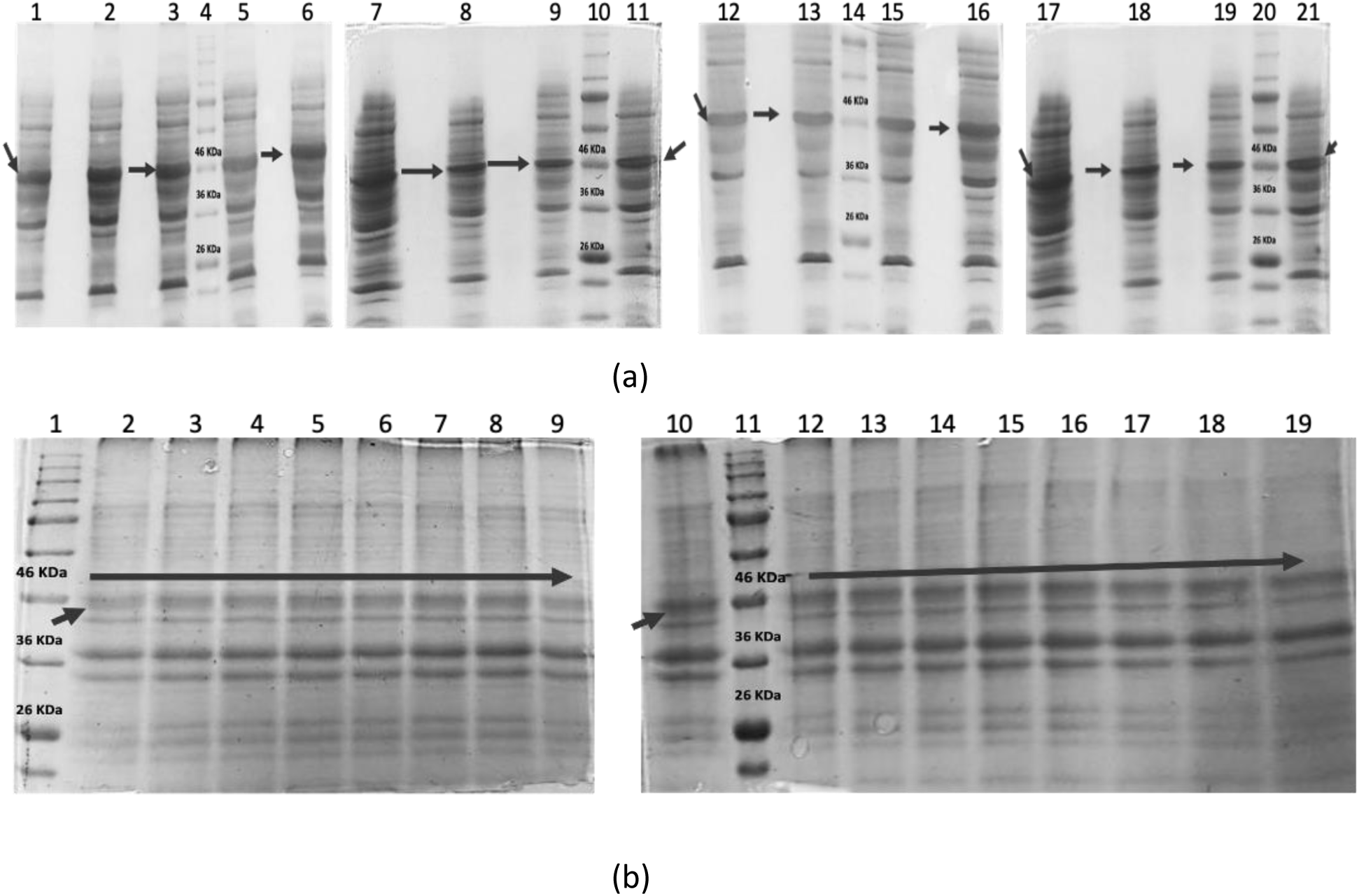
**Recombinant protein over-expression profile of *rsep* metalloprotease WT and mutants expressed in BL21(DE3) *E*. *coli* cells showing the expressed protein bands on 12% SDS-PAGE gel. (a) Soluble protein expression on SDS-PAGE gels, Lane 1,2,3,5,6,7,8,9, *rsep* Mutant M1,M2,M3,M4,M5,M6,M7,M8; Lane 4,10,14,20 Prestained protein ladder; Lane 11,12,13, *rsep* Mutant A1,A2,A3; Lane 15,16,17, *rsep* Mutant G1,G2,G3; Lane 18,19, *rsep* Mutant MG1 , MG2 ; Lane 21, *rsep* Mutant D1. (b) Protein expression in inclusion body fraction on SDS-PAGE gels, Lane 1,11, Prestained protein ladder; Lane 2,3,4,5,6,7,8,9, *rsep* Mutant M1,M2,M3,M4,M5,M6,M7,M8; Lane 10,12,13; *rsep* Mutant A1,A2,A3; Lane 15,16,17; *rsep* Mutant G1,G2,G3; Lane 18,19, *rsep* Mutant MG1, MG2.** M1, M2,M3,M4,M5,M6,M7,M8,A1,A2,A3,G1,G2,G3,MG1,MG2,D1-*rsep* mutant protease. The mutant gene products were named after the modified conditions in the (EP)PCR M1:Mutant developed in 25 µM MnCl_2_, 5.0 mM MgCl_2_; M2:Mutant developed in 50 µM MnCl_2_, 5.0 mM MgCl_2_; M3:Mutant developed in 75 µM MnCl_2_, 5.0 mM MgCl_2_; M4:Mutant developed in 100 µM MnCl_2_,5.0 mM MgCl_2;_ M5:Mutant developed in 25 µM MnCl_2_,10 mM MgCl_2_; M6: Mutant developed in 50 µM MnCl_2_,10 mM MgCl_2_; M7:Mutant developed in 75 µM MnCl_2_, 10 mM MgCl_2_, M8:Mutant developed in 100 µM MnCl_2_,10 mM MgCl_2_; A1:Mutant developed in 0.15 mM dATP, 1.0 mM other dNTPs; A2:Mutant developed in 0.20 mM dATP, 1.0 mM other dNTPs; A3:Mutant developed in 0.25 mM dATP, 1.0 mM other dNTPs; G1:Mutant developed in 0.5 mM dGTP, 0.1 mM other dNTPs; G2:Mutant developed in 0.75 mM dGTP, 0.1 mM other dNTPs; G3:Mutant developed in 1.0 mM dGTP, 0.1 mM other dNTPs; MG1: Mutant developed in 5 mM MgCl_2_; MG2: Mutant developed in 10 mM MgCl_2_; D1:Mutant developed in 5 % v/v dimethylsuphoxide.

The crude enzymes from all the *rsep* mutants and wild recombinant types were compared and analyzed on gelatin-media-agar plates. The hydrolysis zone resulted from the enzyme’s diffusion in the media plates due to the enzyme-substrate reaction between the *rsep* and gelatin. The wild recombinant *rsep* and the mutants showed zones of hydrolysis. In most cases, the hydrolytic zone was comparable to wild recombinant *rsep* except in two cases. In a mutant named ‘A1’, the enhanced proteolytic zone was observed, while in the case of the mutant named ‘G1’, no hydrolysis was visible **(Figure S2)**. Similar experiments were also performed with skim milk agar media plates, which concluded the same results. Many studies described the qualitative estimation of enzyme activity on substrate agar plates for comparative screening. One such study involved the detection of extracellular cellulase and xylanase activity from bacterial isolates where carboxymethyl cellulose and Xylan agar media plates showed hydrolytic zones when exposed to cell-free enzymes of different isolates [39]. Also, a study demonstrated the application of a similar quantitative analysis of protease activity by the casein plate method, where the commercial protease (trypsin) enzyme’s subjection exhibited a measurable hydrolysis zone [40].

Furthermore, to screen the *rsep* mutant with enhanced protease activity, quantitative measurement of the proteolytic activity of these mutants was investigated compared to the wild recombinant *rsep*.

### 3.4. Screening of the rsep mutant with enhanced protease activity

A quantitative protease assay was performed to analyze the *rsep* mutant, which exhibited the highest protease activity towards substrate casein (0.6% w/v). The crude enzymes harvested from different *E*. *coli* hosts were used for activity assay analysis. All the generated mutant proteases showed proteolytic activity against casein **(Table S3)**.

The wild recombinant *rsep* (WT) possessed 14.86 IU/ml, comparable to the activity of mutants M1, M2, M3, M4, M5, M6, M7, M8, A2, G2, G3, and D2 ranged in between 14.85 IU/ml to 14.90 IU/ml. Similarly, the *rsep* mutant activities of A3, D1, and MG2 were calculated to be 15.09 IU/ml, 15.34 IU/ml, and 15.42 IU/ml. These also exhibited approximately the same proteolytic activity as that of WT. The mutant *rsep* MG1 showed an insignificant increase in protease activity (17.15 IU/ml) compared to WT. Only two mutant *rsep,* A1 and G1, showed significantly altered protease activity. The A1 exhibited 7.7 times higher activity than WT, whereas G1 showed a complete activity loss **(Table S3)**. Hence, the A1 *rsep* mutant was selected for further studies. It was purified to homogeneity and studied for the functional and structural modifications describing the improved properties of the mutant protease.

In similarly reported studies, the screening of improved enzymes was performed using the quantitative enzyme activity assay. One recent study involved improving the organic solvent resistance of keratinase BLk by directed evolution. The enzyme activities of the mutant keratinases were checked against casein as a substrate to investigate the improvement in specific activity of the mutant enzymes, which remained the same as the wild-type keratinase. Still, the mutant showed a better half-life towards organic solvents [16]. Another study used a random mutagenesis approach to develop and select solvent-stable haloperoxidase enzymes from the genome of *Streptomyces aureofaciens*. The mutant enzymes were screened using the standard activity assay against the substrate monochlorodimedone (MCD), and highly stable mutant enzymes were screened by quantifying and comparing the residual haloperoxidase activity (MCD reductions) of wild-type and mutant enzymes [41].

### 3.5. Affinity-based purification of the evolved mutant rsep A1

The screened *rsep* A1, cloned in pET22b+ plasmid vector and expressed in *E*. *coli* BL21(DE3), was further purified by Ni-NTA affinity chromatography using a cm X 15 cm column. The bound proteins were eluted by imidazole gradient (150–500 mM in 0.1 M tris HCl buffer; pH 8.0). The eluted proteins were run into a 12 % polyacrylamide gel **(Fig. 3)**. The A1 was eluted at 250 mM imidazole concentration corresponding to a purified protein band of 92 kDa molecular mass. Further, the activity staining confirmed the protease activity in the purified proteases, showing clear proteolytic zones against the background colour of the polyacrylamide gel at the respective molecular size of the proteases, which was calculated to be 50.68 IU/mg as per the activity assay. Further, the activity staining confirmed the protease activity in the purified *rsep* A1 protease, showing clear proteolytic zones against the background colour of the polyacrylamide gel at the respective molecular size of the protease at 92 kDa, showing it to be a dimer as compared to *rsep* (reported to be 46 kDa). The *rsep* protease activity was reported as 6.301 IU/mg as per the activity assay, whereas the mutant *rsep* A1 possessed an activity of 50.68 IU/mg. Hence, the *rsep* gene was evolved through directed evolution to enhance the specific activity of the protease via. a developed mutant *rsep* A1 protease with an improved specific activity 8.04 times the specific activity of recombinant WT *rsep* **(Table S4)**.

**Fig. 3.**
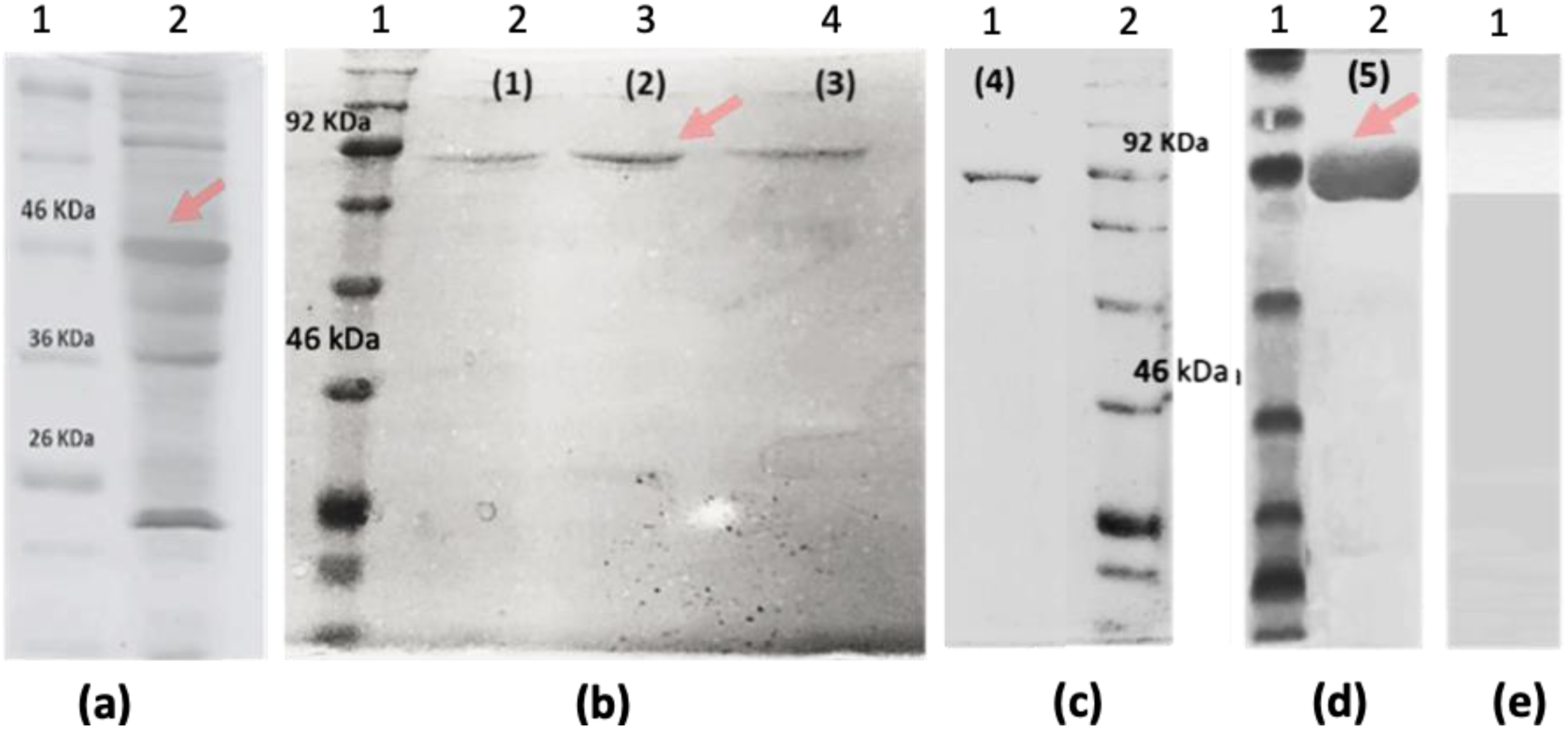
The protein purification profile of mutant *rsep* (a) Reducing SDS-PAGE protein profile of *rsep* mutant metalloprotease (b), (c), and (d) Native PAGE purification profile of *rsep* mutant metalloprotease through Ni-NTA affinity column chromatography showing the purified protein bands on 12% polyacrylamide gels. (a) Lane 1, Prestained protein ladder, Lane 2, Reducing SDS-PAGE protein overexpression profile of *rsep* mutant metalloprotease (b) Lane 1, Prestained protein ladder; Lane 2,3,4, Purified protein fraction (fractions 1,2,3) profiles of 200 mM imidazole based elutes of *rsep* mutant metalloprotease (c) Lane 1, Protein fraction (fraction 4) profile of 200 mM imidazole based elutes of of *rsep* mutant metalloprotease; Lane 2, Prestained protein ladder (d) Lane 1, Prestained protein ladder; Lane 2, Protein fraction profile of 250 mM imidazole based elutes of *rsep* mutant metalloprotease (e) zymography gel showing zone of *rsep* mutant metalloprotease activity on the polyacrylamide gel.

Various attempts have been made to describe the purification of evolved mutant enzymes with improved properties. A similar study exhibited the increase in thermostability of prolyl endopeptidase (PEP) (from the genome of *F*. *meningosepticum*) by error-prone mutagenesis-based directed evolution. The purification of the mutant PEP was performed through CM-cellulose ion exchange chromatography, leading to a considerably thermostable mutant with 51.2 IU/mg specific peptidase activity [17]. Another such study showed the improved catalytic efficiency of the recombinant mutant subtilisin E from the genome of *Thermus aquaticus*, purified by ion exchange technique using the CM-cellulose column. The specific activity was reported to have insignificantly increased from 132 IU/mg to 151 IU/mg in the generated mutant by the site-directed mutagenesis approach. At the same time, the thermostability was considerably improved (with a two times half-life of the mutant compared to wild-type subtilisin at 45 °C and 55 °C temperatures) through the mutational alterations [14]. Another study described the purification of a recombinant Savinase enzyme using Ni-NTA affinity chromatography after the evolution of the enzyme through a microdroplet-based directed evolution approach, leading to higher activity and 5-fold better catalytic rate mutant protease compared to the commercially available protease [13]. Some of the other examples of recombinant proteases from the related genus, especially *Bacillus* sp. such as *Haloalkalobacillus lehensis* [35] and *Bacillus lehensis* [38], have been previously reported to be purified by Ni-NTA chromatography and eluted in the imidazole concentration ranging from 250 mM-500 mM and 200 mM-300 mM, respectively. The activity of purified recombinant proteases from *Bacillus* exhibited an activity of less than 3.0 IU/mg in most of the cases.

### 3.6. Enzymatic and structural properties of purified rsep A1 mutant as compared to wild-type recombinant rsep

#### 3.6.1. pH, temperature optima, and stability

The recombinant *rsep* A1 mutant was characterized to check its enzymatic properties compared to the recombinant *rsep* protease **(WT)** from the genome of *Exiguobacterium* sp. TBG-PICH-001. The pH optima for *rsep* A1 was estimated to be 8.5. This was comparable to the optimum pH value of recombinant *rsep,* reported as 9.0 **(Fig. 4a)**. The temperature optima of *rsep* A1 exhibited a slight shift to 40 °C compared to *rsep*, which was reported to be 35 °C **(Fig. 4b)**. The mutant *rsep* did not lose alkaline stability and was comparably stable to the wild-type exhibiting an enzyme half-life of about 20, 30, 32 h at pH 7, 8.5 and 9.5 **(Fig. 5a)**. The temperature stability of the mutant *rsep* was a little better than the wild-type with an improvement of 1.5x compared to half-life the wild-type at all temperatures (Fig. 5b). Hence, the mutant protease was stable in alkaline pH and warm temperatures. This renders it attractive for detergent applications and useful for proteolysis under alkaline conditions. Interestingly, the mutation did not cause any alteration to the pH-temperature optima and stability of the *rsep* A1 protease, which was equally stable and could be better utilized for industrial applications due to enhanced proteolytic activity. Various microbial alkaline enzyme production and purification studies report the enzyme’s pH and temperature optima calculations and their stability in different pH and temperature ranges. Some of these studies were reported for an alkaline protease from haloalkaliphilic *Bacillus* sp. [27], alkaline lipase and protease from *Pseudomonas aeruginosa* (PseA) [28,42], detergent stable protease from a halophilic *Bacillus* sp. EMB9 [43]. All these studies reported the alkaline pH optima and moderate temperature optima ranging from 35-45 °C. The stability of enzymes from isolated bacteria from different habitats under these studies was stable in the alkaline range with moderate temperature stability. Other studies related to recombinant proteases from the genome of *Bacillus* sp., such as *Haloalkalobacillus lehensis* [35] and *Bacillus lehensis* [38], involving recombinant protease overexpression also described similar trends of protease pH and temperature optima and stability.

**Fig. 4.**
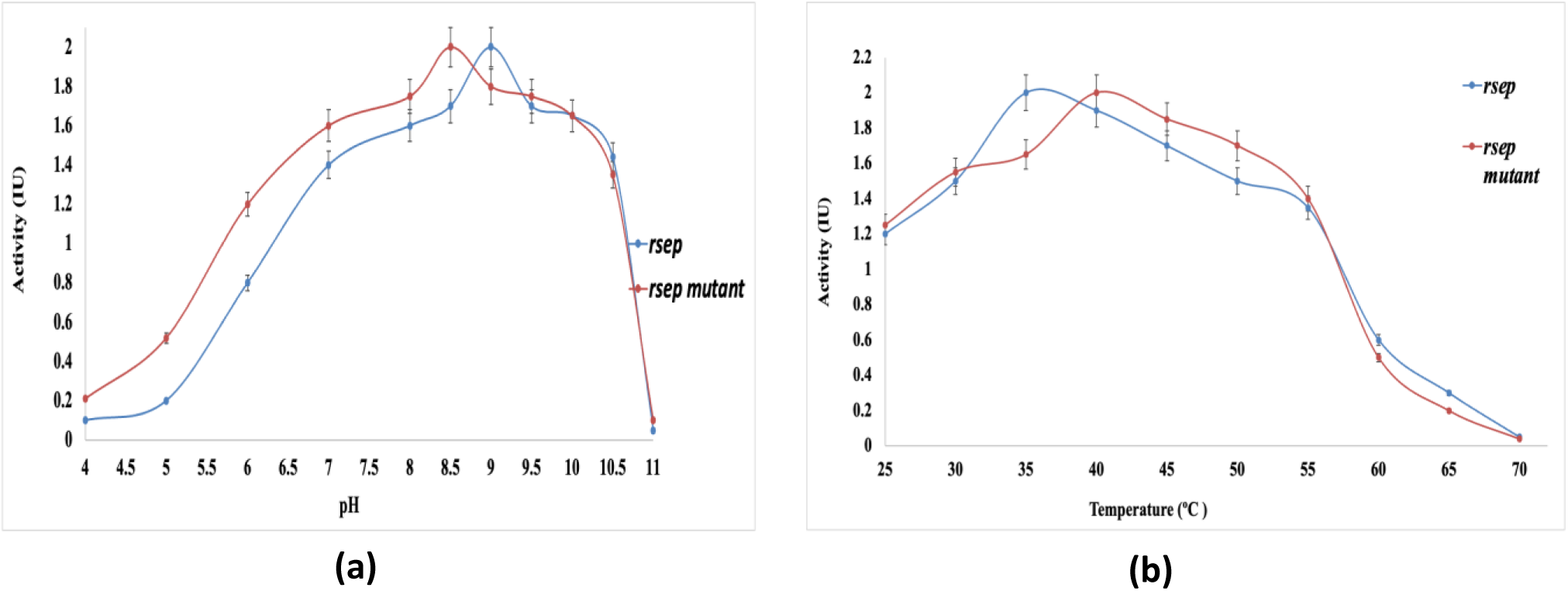
Determination of (a) pH and (b) temperature optima of *rsep* mutant metalloprotease compared to Wild Type (WT)-*rsep* metalloprotease. Values are the mean SD of triplicate determination. The effect of pH on pure enzymes was studied by assaying the enzymes at different pH. Different pH was maintained by dissolving the pure enzyme in buffers of different pH values (0.1M acetate buffer (pH 4.0-6.0); 0.1M phosphate buffer (pH 6.0-8.0); 0.1M Tris-HCl buffer (pH 8.0-9.0), and 0.1M Glycine-NaOH buffer (pH 9-10) was used), at 37 ^◦^C. The temperature optimum for the purified protease was determined by performing the assay at various temperatures. The protease activities were determined under standard assay conditions.

**Fig. 5.**
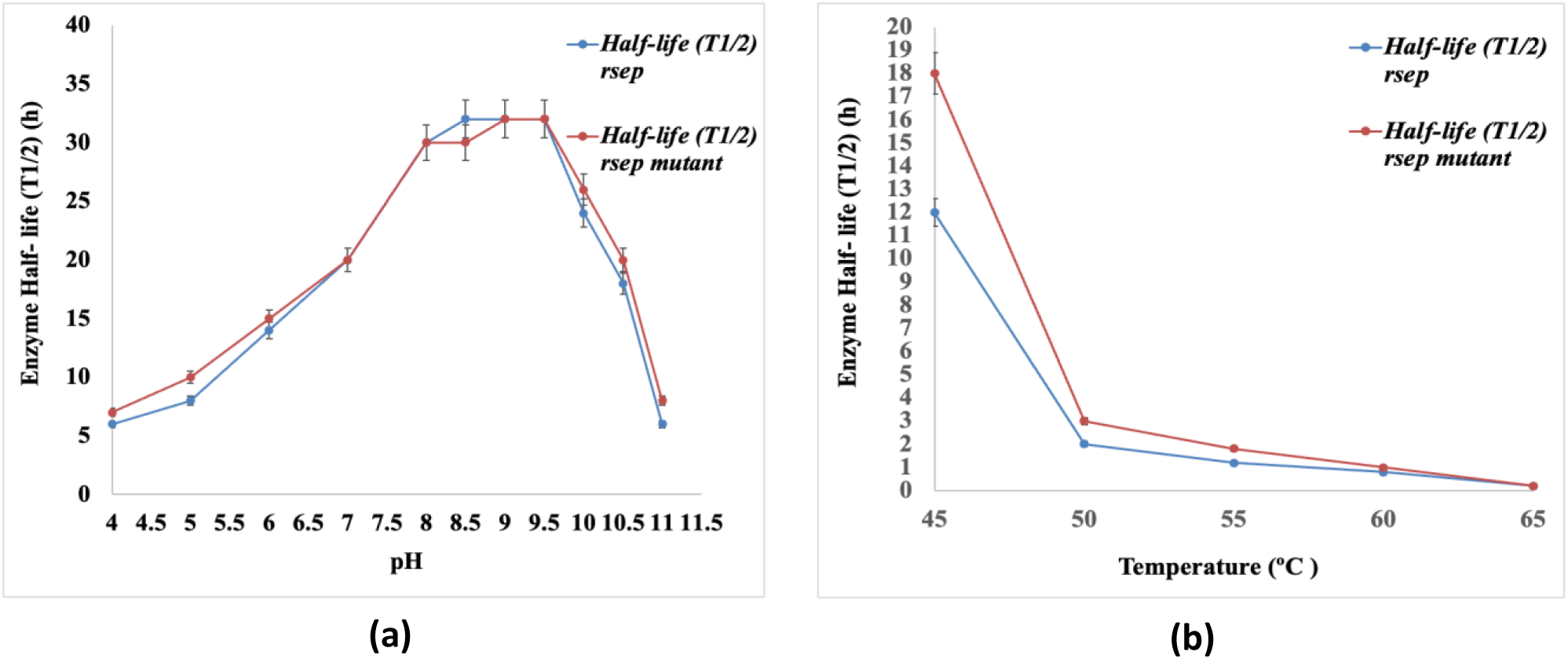
Determination of (a) pH and (b) temperature stability of *rsep* mutant metalloprotease compared to Wild-type (WT)-*rsep* metalloprotease. Values are mean SD of triplicate determinations. The pH stability was studied by pre-incubating the enzymes in buffers of different pH values (0.1M acetate buffer (pH 4.0-6.0); 0.1M phosphate buffer (pH 6.0-8.0); 0.1M Tris-HCl buffer (pH 8.0-9.0), and 0.1M Glycine-NaOH buffer (pH 9-10) was used), at 37 ^◦^C . The thermal stability was studied by incubating the enzymes at different temperatures. The aliquots were withdrawn at different intervals, and the enzyme half-lives were determined by calculating the residual activities under standard assay conditions. The pH conditions for performing the temperature stability experiments were pH optima for the enzyme (pH 8.5).

#### 3.6.2. Substrate specificity

The affinity of the A1 was determined towards various natural and synthetic protein substrates such as Casein, Azocasein, Gelatin, Haemoglobin, Albumin, Ovalbumin, Collagen, Keratin azure, BTEE, ATEE, BAEE, and BAPNA **(Table 1)**. The recombinant WT *rsep* and the mutant *rsep* exhibited the same substrate affinity trend. These demonstrated the highest affinity towards casein > azocasein > gelatin > haemoglobin > albumin > ovalbumin > collagen > keratin. The comparative substrate affinity of mutant *rsep* was enhanced for all the substrates compared to the recombinant WT *rsep*. Briefly, both enzymes preferred natural substrates like Casein, Gelatin, Collagen, and Albumin, but a decreased preference or broader specificity was observed in the mutant *rsep* due to mutational alterations. However, trypsin/chymotrypsin and esterase-like activities were absent in recombinant WT *rsep* and mutant *rsep* A1 metalloproteases. Further, the catalytic parameters viz. K_m_, V_max,_ K_cat_, and Kcat/Km were calculated for both enzymes towards Casein, Azocasein, and Gelatin substrates. Similar substrate preference has been reported in the case of a serine protease from cold-adapted *Planococcus* bacterium, which hydrolysed various proteolytic substrates with the highest affinity towards casein and gelatin, respectively, with a K_m_ of 14.61 mg/mL and V_max_ of 3.104 × 103 μg min^-1^ mg^-1^ against casein as substrate [44]. Similarly, in a study, the heterologous expression was checked for a recombinant protease of an *Aspergillus ochraceous* strain that showed the highest affinity towards a chromogenic substrate (Tos-Gly-Pro-Arg-pNA) with Km and V_max_ value of 435 μM, and 34.4 nmol/min, respectively [45].

#### 3.6.3. Kinetic properties of the proteases

The K_m_, V_max._ for recombinant WT *rsep* and mutant *rsep* A1 metalloproteases towards Casein was calculated to be 391.53 μM, 29.59 μM/ml/min, and 185.87 μM, 30.74 μM/ml/min, respectively which was similar towards Azocasein (K_m WT_ *_rsep_*, V_max WT_ *_rsep_*, 393.4 μM,29.27 μM/ml/min; K_m mutant_ *_rsep_* _A1_, V_max mutant_ *_rsep_* _A1_, 187.6 μM, 31.68 μM/ml/min). The K_m_, V_max._ for recombinant WT *rsep* and mutant *rsep* A1 metalloproteases towards Gelatin was estimated to be 1415.90 μM, 31.32 μM/ml/min, and 1037.20 μM, 32.20 μM/ml/min **(Table 2)**. Briefly, the K_m_ value is decreased for the mutant *rsep* A1 metalloprotease towards all the substrates, demonstrating its better affinity towards proteolytic substrates. The V_m_ value of the mutant remained unaffected compared to the wild recombinant type metalloprotease. The relative catalytic efficiency was also calculated for the recombinant mutant and wild-type *rsep*. The result showed that the relative catalytic efficiency of the mutant was 4.21, 4.08, and 2.84 times that of the recombinant wild-type *rsep* towards casein, Azocasein, and Gelatin, respectively. The calculated increase in the catalytic efficiency suggests a significant improvement in the catalytic efficiency of the evolved mutant through mutagenesis. Several attempts related to enzyme improvement through directed evolution have been attempted where comparative catalytic efficiency of the mutant compared to wild-type recombinant enzyme was described like mutant prolyl endopeptidase (PEP), which showed negligible effect on catalytic parameters like K_m_, V_max_ of mutant PEP (0.1 mM, 200 s^-1^) compared to the wild type (0.084 mM, 210 s^-1^) after two rounds of error-prone PCR [17], a mutant keratinase which showed same K_m_, V_max_ value (0.084 mM) [16]. In general, alkaline proteases from different related bacterial sources have been reported for efficient catalytic parameters; for instance, a detergent protease from a *Bacillus* sp. EMB9 with a K_m_, V_max_ value of 2.22 mg ml^-1^, 1111.11 U ml^−1^ [43], an alkaline protease from haloalkaliphilic *Bacillus* sp. K_m_, V_max_ value of 2.0 mg ml^-1^, 289.0 μg min^−1^ [27], alkaline protease from *Pseudomonas aeruginosa* with K_m_, V_max_ value of 2.7 mg ml^−1^, 3.0 μmol min^−1^, respectively [28]. Some other studies have reported recombinant proteases from *Haloalkalobacillus lehensis* [35] and *Bacillus lehensis* [38] with a K_m_, V_max_ value of 0.2 mg ml^−1,^ 25.0 nmol mg^-1^s^-1^; 1.38 mg ml^−1^, 27.14 μmol min^−1^, respectively.

**Table 2.** Kinetic properties of the *rsep* wild-type and *rsep* mutant metalloproteases

| **Enzyme** | **Substrate** | **K<sub>m</sub>**<br>**(μM)** | **V<sub>m</sub>**<br>**(μM/**<br>**mL/min)** | **K<sub>cat</sub>**<br>**(min<sup>-1</sup>)** | **K<sub>cat</sub>/K<sub>m</sub>**<br>**(μM/min)** | **Relative catalytic**<br>**efficiency**<br>**(Mutant :Wild**<br>**type)** |
| --- | --- | --- | --- | --- | --- | --- |
| **Wild-Type**<br>**(WT)** | **Casein** | 391.5 | 29.59 | 9.433 | 0.024 | **4.21** |
| **Mutant (M)** |  | 185.9 | 30.74 | 18.86 | 0.101 |  |
| **Wild-Type**<br>**(WT)** | **Azocasein** | 393.4 | 29.27 | 9.32 | 0.027 | **4.07** |
| **Mutant (M)** |  | 187.6 | 31.68 | 18.66 | 0.110 |  |
| **Wild-Type**<br>**(WT)** | **Gelatin** | 1415.9 | 31.30 | 9.59 | 0.0067 | **2.84** |
| **Mutant (M)** |  | 1037.2 | 32.19 | 19.75 | 0.0190 |  |

#### 3.6.4. Effect of surfactants, protease inhibitors, and metal ions

The recombinant WT *rsep* and mutant *rsep* A1 metalloproteases were checked for their stability in the presence of detergents and reducing agents, viz. Tween 80, Triton-X-100, S.D.S., Urea, β-Mercaptoethanol, Glutathione, Iodoacetamide, p-Aminobenzidine, and PMSF **(Table 3)**. Both proteases showed high stability in the presence of these reagents with residual activity of 93, 92, 71, 72, 69, 97, 92, 81, and 98%, respectively, indicating them as surfactant stable. Surfactant stability is mandatory for the alkaline proteases used as detergent additives. The *rsep* A1 did show a lesser residual protease activity in the presence of DTT, a reducing agent (23%) and 1,10-Phenanthroline, a metalloprotease inhibitor (10%). These results were similar to the stability properties of *rsep* (WT). Hence, the recombinant mutant *rsep* A1 did not lose its stability towards surfactants, and it is quite stable in the presence of inhibitors. Hence, it could be considered a potential and better candidate for applications in detergent formulations than wild recombinant *rsep*. The effect of metal ions (Zn^2+^,Ca^2+^,Mg^2+^,Pb^+^,Ni^2+^,Cu^2+^,Fe^2+^, and K^+^) on the protease activity of mutant *rsep* A1 metalloprotease was checked. The residual activity of the mutant remained unaltered (100%) in the presence of Mg^2+^, Fe^2+^, and K^+,^ while the activity increased in the presence of Zn^2+^ and Ca^2+^, which was 240% and 115%, respectively. A decreased residual activity was shown in the presence of Pb+, Ni^2+^, and Cu^2+^, which were 65%,48%, and 24%, respectively. These results corresponded similarly to the effect of metal ions on the protease activity of wild recombinant *rsep* except for the effect of Zn^2+^, where the residual protease activity for the mutant improved to 413% (4.13 times to that of the control ie.100%). In the presence of Zn^2+^, the *rsep* (WT) exhibited an improved activity, which was 2.4 times compared to the control. Hence, after the mutation, the evolved *rsep* A1 has increased interaction with Zn^2+^ ions, probably because of the increase in the polar amino acid residues interacting with zinc ions and enhancing the substrate proximity with the active site of the enzyme *rsep*A1. Similarly, the recombinant protease obtained from *Bacillus lehensis* JO-26 was stimulated in the presence of Ca^2+^ ions (5mM), and SDS (1% w/v) [35]. Another such study involved the heterologous expression of a serine protease of *Bacillus velezensis* SW5 in the *B*. *subtilis,* which was found stable in the presence of all the metal ions (Ca^2+^, Mg^2+^, Co^2+^, Cu^2+^, Zn^2+^, Mn^2+^, Fe^2+^) except Ni^+^, and Hg^2+^ [46].

**Table 3.** Effect of protease inhibitor, surfactants, and metal ions on the purified *rsep* mutant metalloprotease

| **Inhibitor/surfactants/**<br>**metal ions** | **Conc.** | **WT relative**<br>**activity (%)** | **Mutant relative**<br>**activity (%)** |
| --- | --- | --- | --- |
| **None** | **-** | **100** | **100** |
| **Tween 80** | **0.2 %** | **93± 2** | **93± 1** |
| **Triton- X 100** | **0.2 %** | **92± 1** | **92± 1** |
| **SDS** | **0.2 %** | **71± 0.5** | **71± 0.5** |
| **Urea** | **0.2 %** | **72± 2** | **72± 2** |
| **β- mercaptoethanol** | **0.2 %** | **69± 0.5** | **21± 1** |
| **Glutathione** | **5 mM** | **97± 0.5** | **97± 0.5** |
| **DTT** | **5 mM** | **23± 0.7** | **23± 2** |
| **Iodoacetamide** | **5 mM** | **92± 2** | **92± 3** |
| **p-aminobenzidine** | **5 mM** | **7± 1** | **81± 0.5** |
| **PMSF** | **5 mM** | **100± 1** | **98± 2** |
| **1,10-Phenanthroline** | **10 mM** | **12± 0.5** | **10± 2** |
| **EDTA** | **10 mM** | **2± 0.1** | **2± 0.5** |
| **Zn<sup>2+</sup>** | **2 mM** | **240± 2** | **413± 4** |
| **Ca<sup>2+</sup>** | **2 mM** | **120± 2** | **100± 3** |
| **Mg<sup>2+</sup>** | **2 mM** | **100± 0.5** | **100± 0.5** |
| **Pb<sup>2+</sup>** | **2 mM** | **63± 0.2** | **95± 0.5** |
| **Ni<sup>2+</sup>** | **2 mM** | **49± 2** | **92± 1** |
| **Cu<sup>2+</sup>** | **2 mM** | **22± 0.6** | **93± 0.5** |
| **Fe<sup>2+</sup>** | **2 mM** | **96± 2** | **100± 2** |
| **K<sup>+</sup>** | **2 mM** | **94± 0.5** | **94± 1** |

#### 3.6.5. Organic solvent-stability

The exciting feature of our native *Exiguobacterium* isolate’s proteases was their stable proteolytic property in the non-aqueous system. The recombinant *rsep* also showed relatively stable in the range of organic solvents. Hence, it was necessary to investigate whether the mutational alteration caused any decrease or increase in the stability of the mutant *rsep* A1 towards various organic solvents **(Fig. 6)**. The results of organic solvent stability of the mutant *rsep* A1 were comparable to that of the recombinant WT *rsep*. They retained 100 % activity in non-polar solvents like tetradecane, dodecane, and decane. After that, they start losing activity as a function of decreasing log P value (The partition-coefficient P in an n–octanol–water system, where P is the ratio of the solubility of the solvent in n-octanol to its solubility in water). As per the established principles, protease loses its activity quickly in polar solvents. Since polar solvents strip the essential water molecules from the enzyme’s active site, such activity loss was understandable [47,48]. Hence, the mutant protease did not lose solvent-stable properties and can be applied more inexpensively for industrial applications such as laundry additives and reverse biocatalysis (peptide synthesis) compared to the recombinant *rsep* WT. Similar observations have been made for the recombinant protease from *Bacillus lehensis* JO, which exhibited high stability (solvent conc. 10% v/v) in the presence of n-hexane and loss in the DMF, propanol, chloroform, glycerol, toluene, n-butanol, benzene, methanol, and ethanol [35]. A recombinant serine protease from *Fusarium graminearum* exhibited similar features of higher stability in hydrophobic solvents like hexane, dodecane, and tetradecane while lower stability in the presence of polar solvents like ethanol, butanol, propanol, chloroform, etc. [49].

**Fig. 6.**
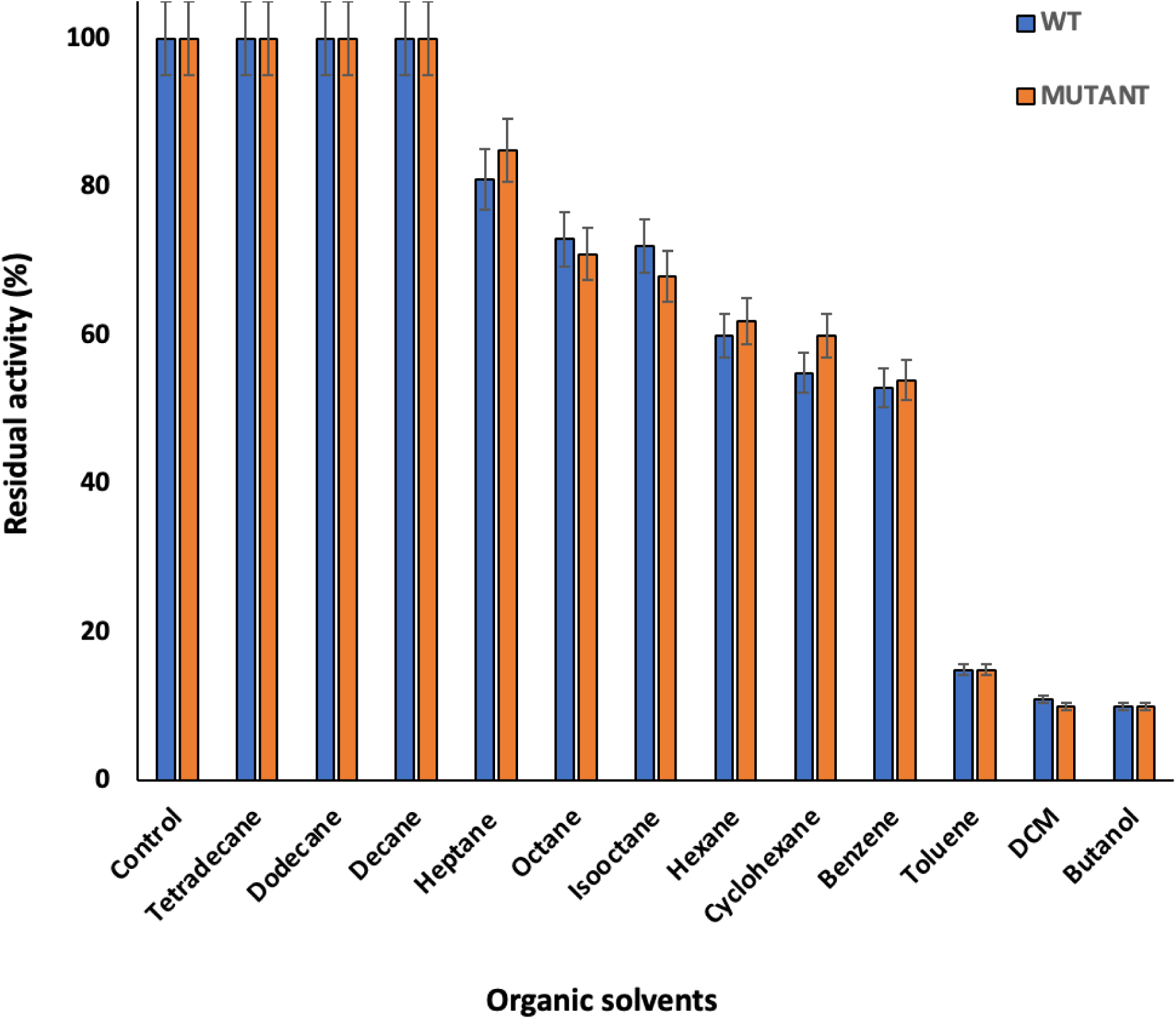
Estimation of the organic solvent stability of mutant *rsep* metalloprotease in various organic solvents compared to Wild type(WT)-*rsep* metalloprotease. A known amount of purified proteases were dissolved in Tris-HCl buffer (0.1M; pH 9.0 and 8.5 for wild type(WT) and mutant *rsep* metalloprotease respectively). One mL of each enzyme was separately mixed with 3 mL of various organic solvents. The mixture was sealed in parafilm and incubated at 200 rpm with constant shaking at 37 ^◦^C for 48 h. The organic solvent untreated sample remains the control, kept under similar conditions. The samples were checked for residual protease activity as per the standard assay conditions. Values are mean ±SD of triplicate determination.

#### 3.6.6. Biophysical studies and their correlation with the primary structure of the rsep mutant A1

Comparative biophysical studies were performed for *rsep* WT and *rsep* mutant through Fluorescence and Circular Dichroism (CD) spectroscopy, and the melting temperature was also compared for both the proteases by Differential Scanning Calorimetry (DSC). The fluorescence spectra showed a decreased fluorescence intensity of *rsep* A1 compared to *rsep* WT. This inferred the appearance of new polar residues or replacement of aromatic amino acid residues or both in the primary structure of the mutant *rsep* A1, probably due to point mutations generated by directed evolution. The increase in the protease surface polarity is indicative of the fact that the polar amino acid residues (aspartate, glutamate, and histidine) may have quenched the aromatic residues, leading to a decrease in the fluorescence intensity recorded in the fluorescence spectra **(Fig. 7)**. A recent study described the decrease in the fluorescence intensity due to the decrease in the overall surface hydrophobicity of an enzyme and vice-versa after performing a rational surface charge engineering approach to enhance the solvent stability in the dehalogenase enzyme (DHA) [50]. Another study revealed a decrease in the fluorescence intensity of protein due to nearby polar histidine or cysteine residues [51].

**Fig. 7.**
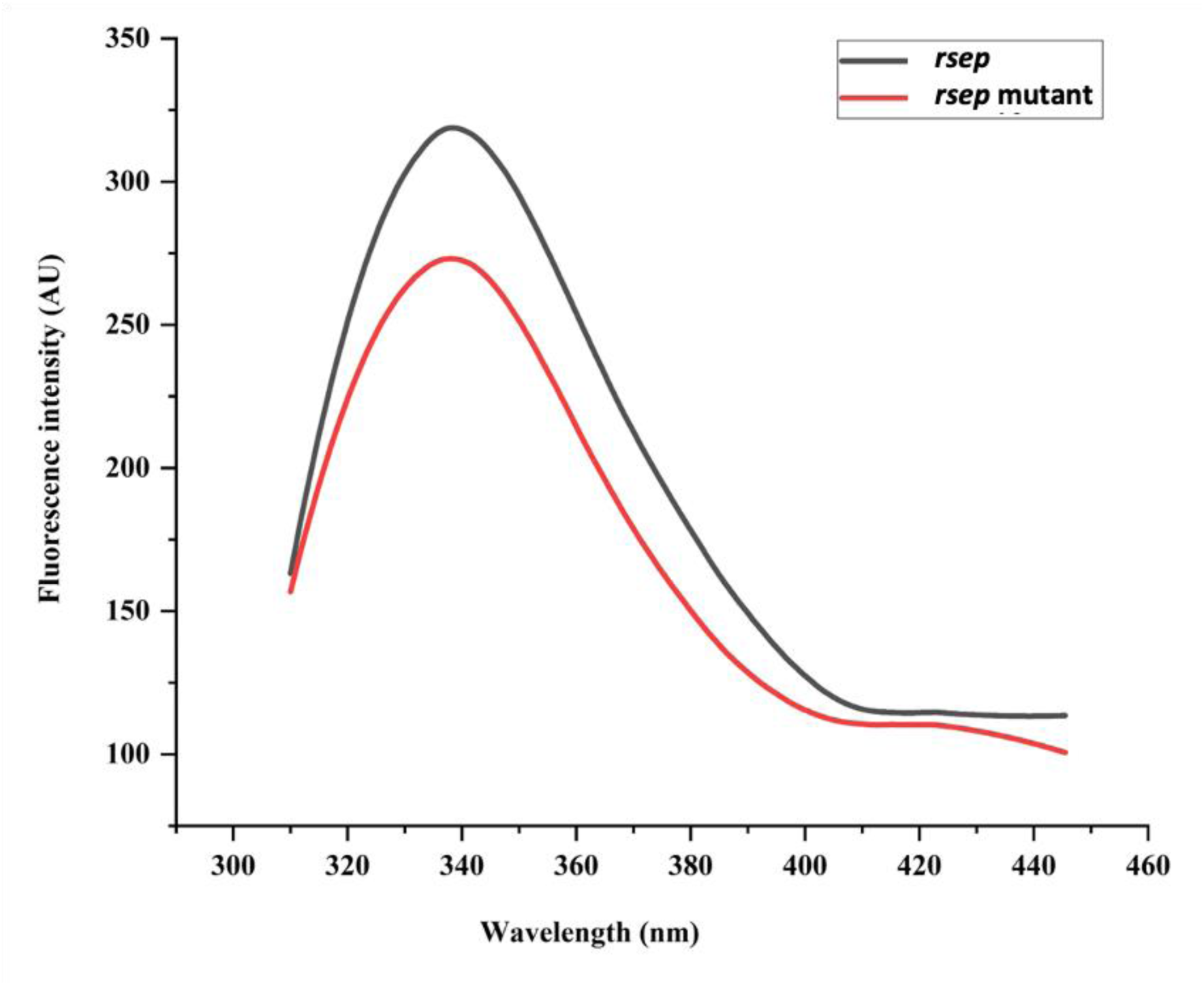
Determination and comparison of tertiary structure of mutant *rsep* metalloprotease and wild-type (WT)-*rsep* metalloprotease. Tertiary structure was determined through fluorescence spectra studies. A known amount of the purified mutant and wild-type (WT)-*rsep* metalloprotease were dissolved in 20 mM Tris-HCl (pH 8.5), and the analyses were performed.

The CD spectra concluded the decrement in the percentage of α-helices compared to *rsep* WT, which indicated substitution of alanine residues, which has the highest propensity of forming alpha helices in the primary structure of mutant *rsep* A1**(Fig. 8)**. The 2D-structure for the mutant *rsep* A1 and Wild-type (WT)-*rsep* were also predicted through I-TASSER online software, which corresponded to decreased α-helix percentage and verified the results obtained from CD data **(Figure S3)**.

**Fig. 8.**
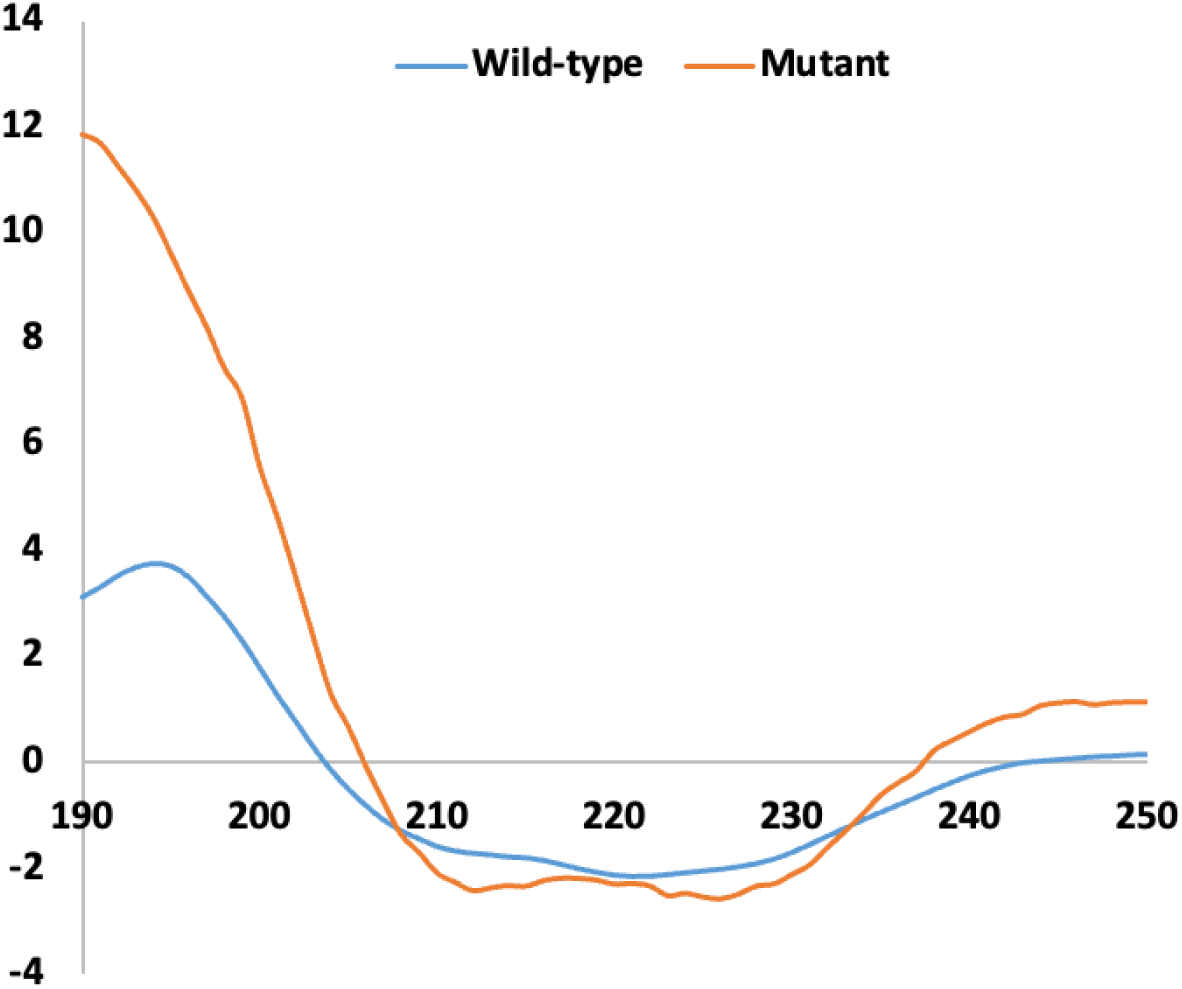
Determination and comparison of secondary structure of mutant *rsep* metalloprotease and wild-type (WT)-*rsep* metalloprotease. (a) The secondary structure was determined through circular dichroism (CD) spectra studies. A known amount of the purified mutant and wild-type (WT)-*rsep* metalloprotease were dissolved in 20 mM Tris-HCl (pH 8.5)

Additionally, the melting temperature for both the proteases was calculated and compared by DSC. The results concluded a shift of 7.31°C in the melting temperature of mutant *rsep* A1 (70.43 °C) compared to *rsep* WT (63.12 °C). This result correlated with the slight increase in the temperature optima of the mutant *rsep* A1 compared to the WT *rsep* **(Figure S4)**.

The nucleotide sequence was identified through cloned mutant plasmid sequencing to investigate the occurrence of point mutations in the primary structure of the mutant *rsep* A1 **(Fig. 9)**. The comparative primary structures (amino acid sequence) obtained using the Expasy Translate online bioinformatic tool were aligned through the Clustal W alignment tool, which clearly showed the appearance of new polar amino acid residues in the primary sequence of the mutant *rsep* A1 compared to the WT *rsep*. These amino acids were chiefly Histidine, Lysine, Glutamic acid, and Aspartic acid. The data presented the highest increase in the number of Histidine residues participating in the active site of such metalloproteases besides probably making the surface more polar and resulting in better interaction with the substrate, increasing its affinity towards the substrate and possibly causing an increased interaction with the Zinc ions to bring the substrate in close vicinity of the enzyme’s active site. These results are well correlated with the fluorescence and CD spectra studies, which inferred the appearance of polar residues like Histidine, Aspartate, etc., and the substitution of Alanine residues corresponding to the increase in the polarity of the enzyme surface and decrease in the overall α-helix percentage of the mutant enzyme, respectively. Hence, the alterations in the predicted primary amino acid structure corresponded to the experimental data of the *rsep* A1 mutant and provided a satisfactory explanation for the increased substrate affinity, relative proteolytic activity, and catalytic efficiency of the evolved *rsep* A1 metalloprotease generated by the directed evolution of the WT *rsep* metalloprotease.

**Fig. 9.**
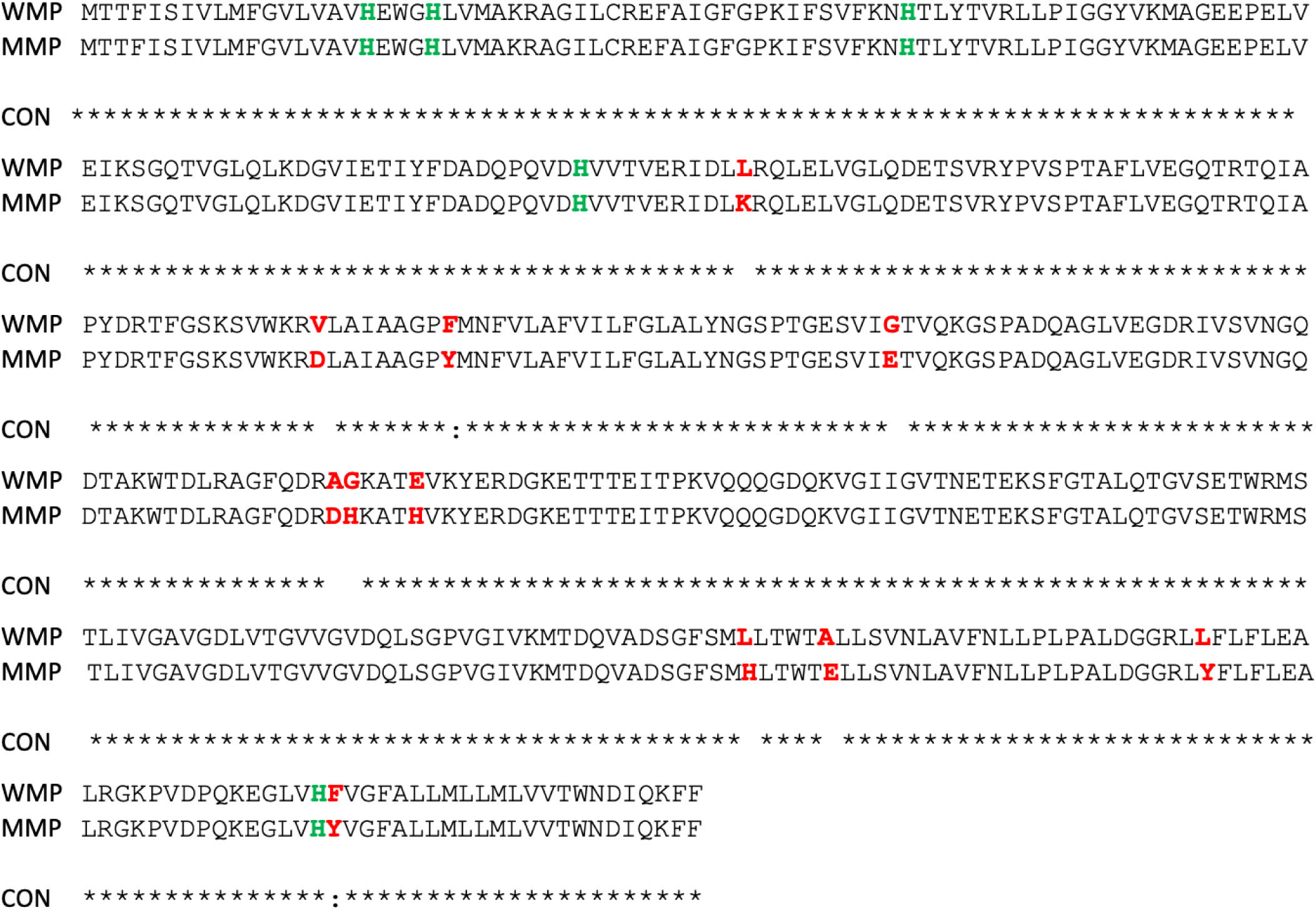
Analysis and comparison of the primary structure of the wild-type metalloprotease (WMP) and mutant *rsep* metalloprotease through sequencing. The modified residues are highlighted in red.

#### 3.6.7. Inductively coupled plasma mass spectrometry (ICP-MS) analysis

The results of ICP-MS analysis showed a clear comparison of the uptake of zinc ions (Zn^2+^) wild-type and mutant *rsep* metalloproteases. The concentration of the ions attained from wild-type and mutant *rsep* were calculated as 33.06 and 100.15 ppb, respectively. This indicated 3.03 times more uptake of zinc ions in the case of the mutant compared to the recombinant wild-type *rsep,* which correlates with the results of sequencing, effects of Zn^2+^ metal ions, and fluorescence data that described the increment in the histidine residues that interact with zinc ions in such metalloproteases which resulted in enhanced protease activity and affinity of the evolved mutant *rsep* metalloprotease through directed evolution. A study showed increased uptake of zinc ions by an IMP-1 metallo-ß-lactamase enzyme when the protein was supplied with excess zinc ions during the growth of *E*. *coli* in the culture media broth. The concentration of the metal ions was assessed through ICP-MS analysis [52]. Another study related to the membrane proteins and metalloproteases of *Bacillus cereus* and *Pseudomonas aeruginosa* for the assessment of protein binding to zinc ions was performed through ICP-MS, which showed zinc to be a significant metal ion factor for rendering the activity of proteins intact [53].

#### 3.6.8. Comparative analysis of the rsep wild-type and rsepA1 mutant metalloprotease

The properties of the wild-type and mutant *rsep* metalloprotease are compared and presented in a table to understand the details of the improvement in the recombinant mutant *rsep*A1 with suitable inferences justifying the structure-function correlation **(Table S5).** Further, the improved properties of wild-type *rsep* v/s mutant *rsep* metalloprotease have also been discussed in another table **(Table 4)**. This table also compares the mutant improvement data with literature that showed improved mutant properties developed through similar enzyme engineering studies.

**Table 4.** Improvement in the properties of mutant v/s wild type *rsep* metalloprotease

| Characteristics | Improvement in *rsep*-mutant compared to wild-type (Current study) | Improvement in BLk-mutant protease (Zhu et al., 2023) | Improvement in EP-B2-mutant endopeptidase (Osorio et al, 2020) | Improvement in PEP-407 mutant endopeptidase (Uchiyama et al, 2000) |
| --- | --- | --- | --- | --- |
| **Thermal stability**<br>[T<sub>1/2</sub> (50 °C)] | 1.5 x | 2 x | 3.5 x | 5.12 x |
| **Solvent stability** | Unaltered | 4 x | - | - |
| **Specific Activity** | 8 x | unaltered | 3 x | 1.1 x |
| **Catalytic efficiency**<br>(K<sub>cat</sub>/K<sub>m</sub>) | 4.2 x | unaltered | 1.6 x | 1.4 x |
| **Substrate specificity** | Broader | Unaltered | Unaltered | Unaltered |
| **Substrate affinity**<br>(K<sub>m</sub>) | x/2.1 | unaltered | x/2.4 | x/1.15 |
| **Melting temperature** | 1.12 x | - | - | - |
| **Zinc metal ion uptake** | 3.03 x | - | - | - |
| **Primary structure (base substitution)** | A-H, G-H | A-Q, D-A | R-I, E-W | G-C, A-C |
| **Secondary Structure**<br>(CD spectra) | Increased alpha character in the mutant protease | Increased alpha character in the mutant endopeptidase | - | - |

## 4. Conclusions

A directed evolution strategy involving the EP-PCR approach could improve recombinant protease properties and enhance their suitability for industrial applications. The current study undertook a recombinant metalloprotease’s directed evolution, characterization, and structure elucidation in this context. A single-round error-prone mutagenesis approach has been employed to develop a mutant metalloprotease with enhanced enzymatic potential, which resulted in a mutant metalloprotease (*rsep* A1) that showed an approximately 8-fold and 4-fold increase in protease activity and catalytic efficiency, respectively. It was purified to homogeneity with a molecular size of 92 kDa (a homodimer). The structural elucidation studies through bioinformatic tools, sequencing data, and comparative biophysical studies of the wild and mutant metalloprotease showed the presence of an increased amount of histidine residues in the mutant metalloprotease besides the appearance of acidic residues like glutamic, aspartic acid, etc. This led to a better interaction between the substrates and the mutant enzyme, which enhanced its affinity and catalytic efficiency. Such evolved proteases could be further studied for in-detail structural information. They may be applied to better tailor such hydrolase groups of enzymes to specific industrial endeavours by incorporating combinatorial enzyme engineering methods.

## Credit authorship contribution statement

**N. Srivastava**: Writing-original draft, Conceptualization, Formal analysis. **S. K. Khare:** Conceptualization, Supervision, Funding acquisition, Editing, and Review.

## Author Statement

The authors declare that they have no conflict of interest and mutually agree to the manuscript submission.

## Acknowledgements

The authors gratefully acknowledge the financial grant provided by the Department of Biotechnology, India (BT/PR26709/AAQ/3/889/2017) for carrying out this study. Nitin Srivastava is grateful to the DST-SERB (Govt. Of India) for the Research Fellowship.

