## Supplementary data file for "Improvement in the enzymatic efficiency of a zinc metalloprotease through random mutagenesis approach: Studies on the enzymatic and structural properties of the mutant"

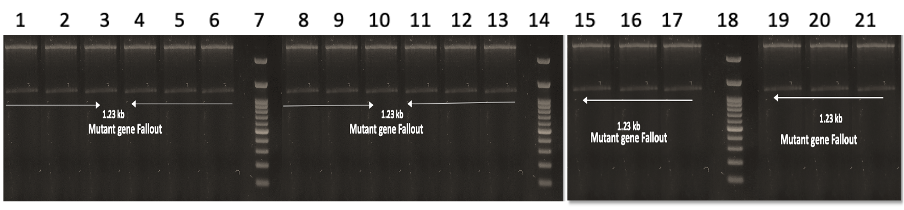

**Figure S1	Clone confirmation of the *rsep* gene-specific fragments (1.23 Kb).** Ethidium bromide-stained gel shows the presence of 1.23 kb fragments of *rsep* when the cloned WT and Mutant pET 22b+ plasmids were observed after digestion with Bam HI and Xho I restriction enzymes. Lane 1,1kb DNA Ladder; Lane 2, *rsep* WT gene fallout; Lane 3-10, *rsep* Mutant M1,M2,M3,M4,M5,M6,M7,M8 gene fallout; Lane 11,12,13, *rsep* Mutant A1,A2,A3 gene fallout, respectively; Lane 14,15,16, *rsep* Mutant G1,G2,G3, respectively; Lane 17,18, *rsep* Mutant MG1 , MG2 gene fallout; Lane 19,20, *rsep* Mutant D1,D2, *rsep* gene fallout.

WT-Wild type *rsep* metalloprotease gene

M1, M2,M3,M4,M5,M6,M7,M8,A1,A2,A3,G1,G2,G3,MG1,MG2,D1,D2-(EP)PCR mutant genes.

The mutant gene products were named after the modified conditions in the (EP)PCR M1:Mutant developed in 25 µM MnCl_2_, 5.0 mM MgCl_2_; M2:Mutant developed in 50 µM MnCl_2_, 5.0 mM MgCl_2_; M3:Mutant developed in 75 µM MnCl_2_, 5.0 mM MgCl_2_; M4:Mutant developed in 100 µM MnCl_2_,5.0 mM MgCl_2;_ M5:Mutant developed in 25 µM MnCl_2_,10 mM MgCl_2_; M6: Mutant developed in 50 µM MnCl_2_,10 mM MgCl_2_; M7:Mutant developed in 75 µM MnCl_2_, 10 mM MgCl_2_, M8:Mutant developed in 100 µM MnCl_2_,10 mM MgCl_2_; A1:Mutant developed in 0.15 mM dATP, 1.0 mM other dNTPs; A2:Mutant developed in 0.20 mM dATP, 1.0 mM other dNTPs; A3:Mutant developed in 0.25 mM dATP, 1.0 mM other dNTPs; G1:Mutant developed in 0.5 mM dGTP, 0.1 mM other dNTPs; G2:Mutant developed in 0.75 mM dGTP, 0.1 mM other dNTPs; G3:Mutant developed in 1.0 mM dGTP, 0.1 mM other dNTPs; MG1: Mutant developed in 5 mM MgCl_2_; MG2: Mutant developed in 10 mM MgCl_2_; D1:Mutant developed in 5 % v/v dimethylsuphoxide; D2:Mutant developed in 10 % v/v dimethylsulphoxide.

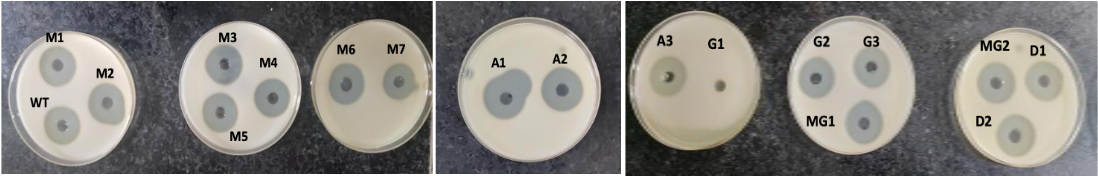

**Figure S2 Comparative Qualitative analysis of the wild-type and mutant *rsep* metalloprotease on NB-gelatin media plates.** The nutrient broth, agar-agar, and gelatin (2% w/v) were used, and media agar plates were constituted. The media plates were bored to make plugs, and the soluble protein fractions (containing the expressed recombinant proteases) were poured and sealed with agar to form agar plugs in the plates. These plates were incubated at 37 °C overnight, and different samples were analysed for the hydrolytic zones by flooding with Frazier’s reagent (15 % HgCl_2_ in 2 N HCl).

M1, M2,M3,M4,M5,M6,M7,M8,A1,A2,A3,G1,G2,G3,MG1,MG2,D1-*rsep* mutant protease. The mutant gene products were named after the modified conditions in the (EP)PCR M1:Mutant developed in 25 µM MnCl_2_, 5.0 mM MgCl_2_; M2:Mutant developed in 50 µM MnCl_2_, 5.0 mM MgCl_2_; M3:Mutant developed in 75 µM MnCl_2_, 5.0 mM MgCl_2_; M4:Mutant developed in 100 µM MnCl_2_,5.0 mM MgCl_2;_ M5:Mutant developed in 25 µM MnCl_2_,10 mM MgCl_2_; M6: Mutant developed in 50 µM MnCl_2_,10 mM MgCl_2_; M7:Mutant developed in 75 µM MnCl_2_, 10 mM MgCl_2_, M8:Mutant developed in 100 µM MnCl_2_,10 mM MgCl_2_; A1:Mutant developed in 0.15 mM dATP, 1.0 mM other dNTPs; A2:Mutant developed in 0.20 mM dATP, 1.0 mM other dNTPs; A3:Mutant developed in 0.25 mM dATP, 1.0 mM other dNTPs; G1:Mutant developed in 0.5 mM dGTP, 0.1 mM other dNTPs; G2:Mutant developed in 0.75 mM dGTP, 0.1 mM other dNTPs; G3:Mutant developed in 1.0 mM dGTP, 0.1 mM other dNTPs; MG1: Mutant developed in 5 mM MgCl_2_; MG2: Mutant developed in 10 mM MgCl_2_; D1:Mutant developed in 5 % v/v dimethylsuphoxide.

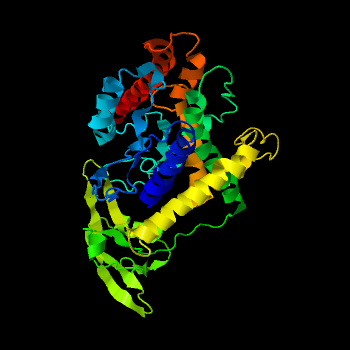

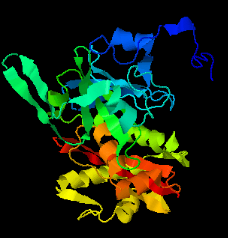

**(ii)**

**(i)**

**Figure S3	I-tasser (online software) based secondary structure prediction of (i) Wild-type (WT)-*rsep* and mutant *rsep* metalloproteases.**

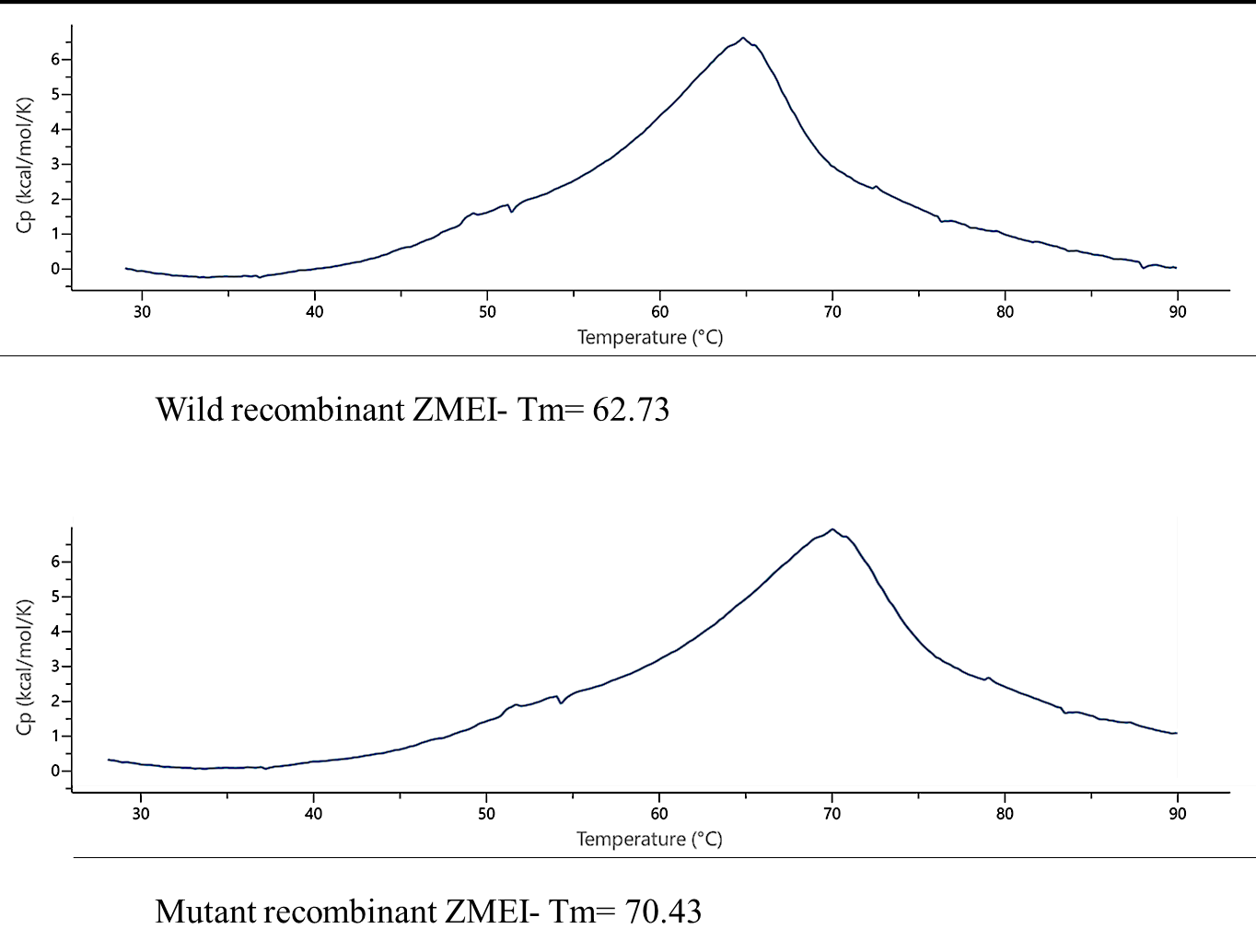

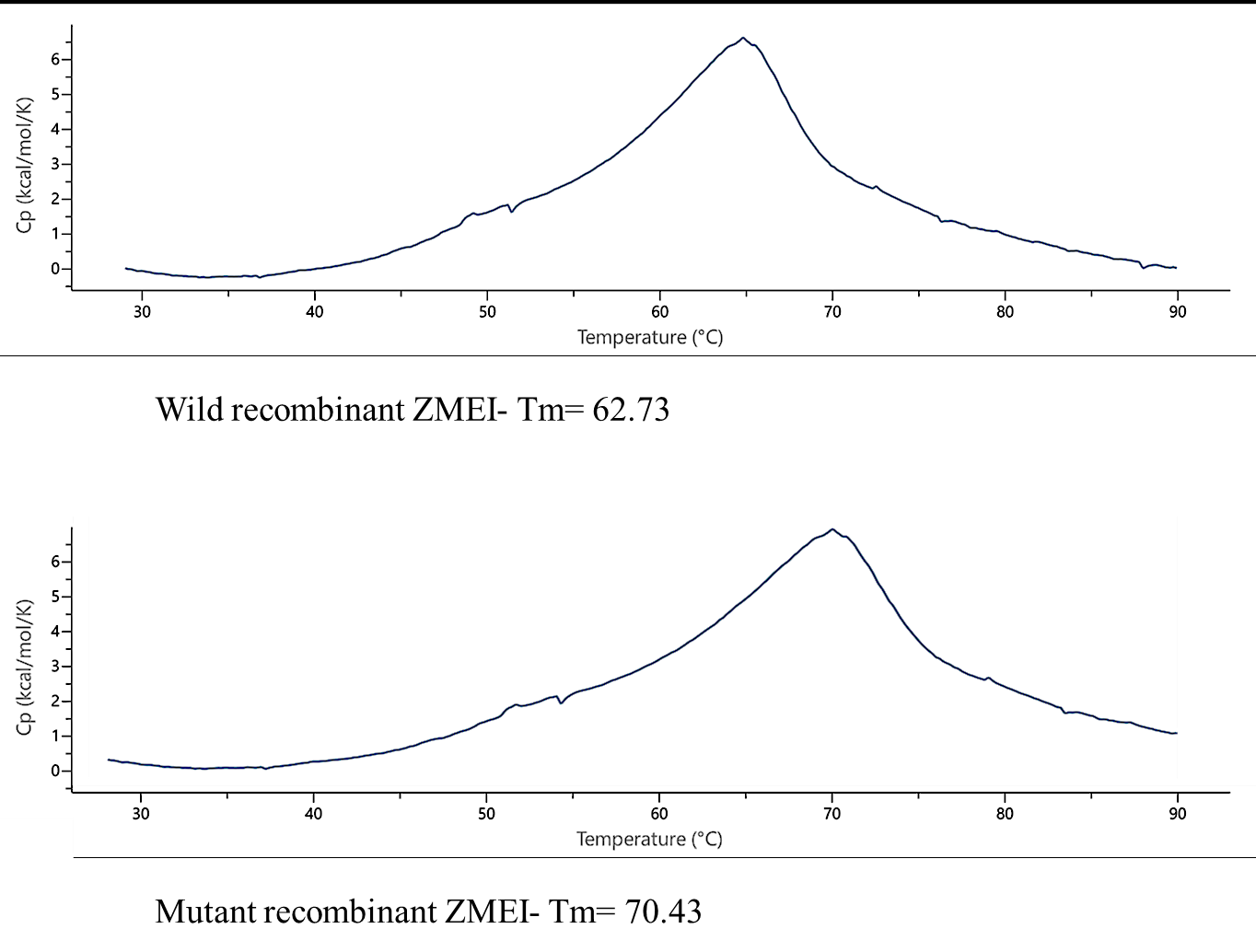

**(a)**

**(b)**

**Figure S4	Determination and comparison of the melting point of (a) wild-type (WT)- *rsep* metalloprotease and (b) mutant *rsep* metalloprotease. The m**elting point was determined through differential scanning calorimetric (DSC) studies. A known amount of the purified mutant and wild-type (WT)-*rsep* metalloprotease was dissolved in 20 mM Tris-HCl (pH 8.5) and used for melting point determination.

**Table S1 Mutant library generation through error prone-polymerase chain reaction (EP-PCR)**

| **S. No.** | **Reagents** | **Basic Conc.** | **Modified Conc.** |
| --- | --- | --- | --- |
| 1 | Tris HCl (pH 8.3) | 10 mM | 10 mM |
| 2 | DMSO | - | 5, 10 % (v/v) |
| 4 | MnCl_2_ | - | 25, 50, 75, 100 µM  with 5 and 10 mM MgCl_2_ |
| 5 | KCl | 50 mM | 50 mM |
| 6 | MgCl_2_ | 1.5 mM | 5, 10 mM |
| 7 | BSA | 0.01% (w/v) | 0.01% (w/v) |
| 8 | dNTPs | 0.2 mM | 1. dGTP- 0.5, 0.75,1.0 mM and Other dNTPs -0.1 mM  2. dATP-0.15, 0.20, 0.25 mM and Other dNTPs -1.0 mM |
| 9 | Primers | 1 µM | 1 µM |
| 10 | Cloned plasmid | 10 ng/ml | 10 ng/ml |
| 11 | Taq DNA polymerase | 25 U/ml | 25 U/ml |

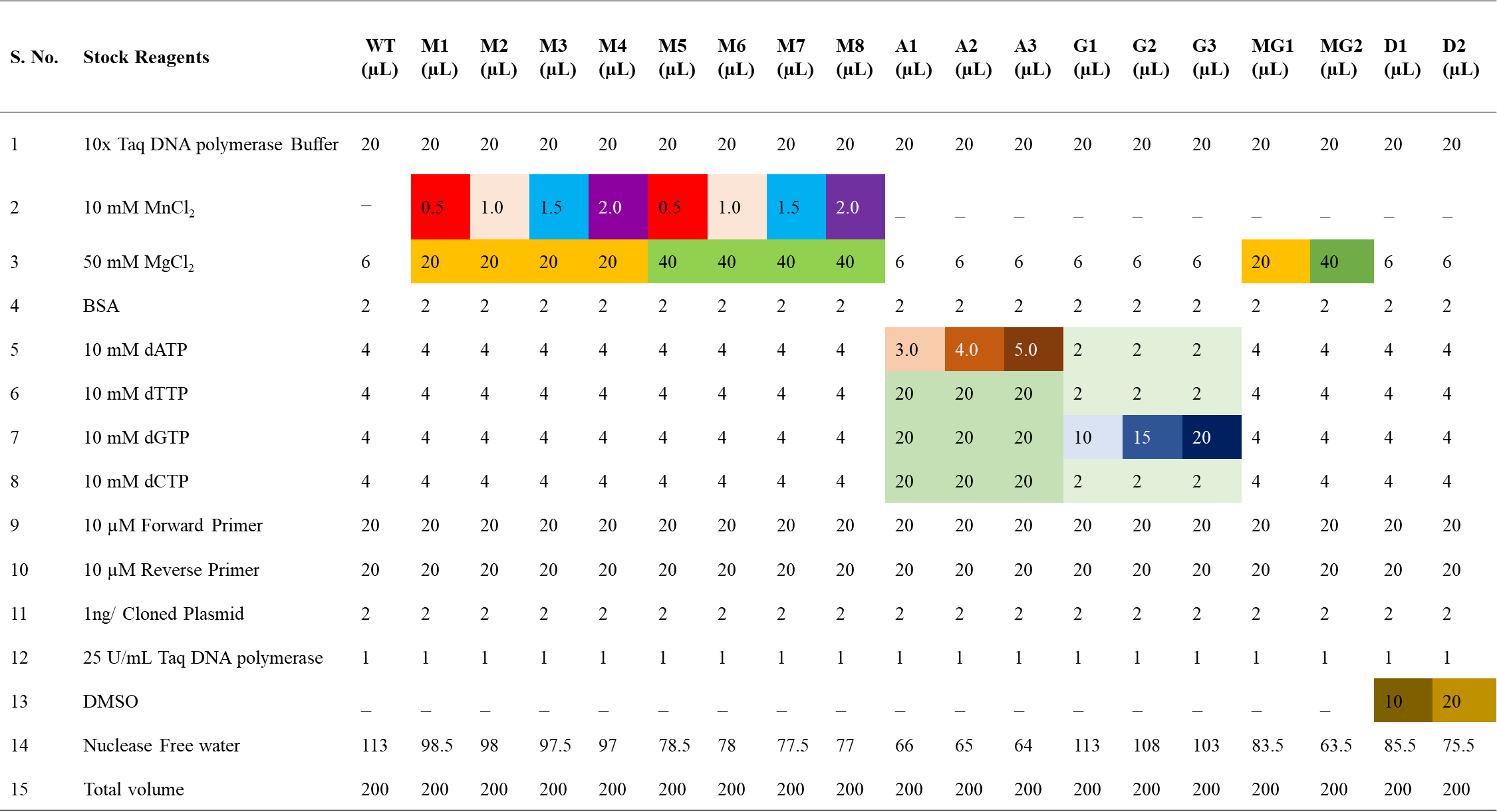
**Table S2	EP-PCR reaction chart for mutant gene library generation**

WT, wild-type; M1-M4, Mutant MnCl_2_- 25, 50, 75, 100 µM with 5 mM MgCl_2_ ; M5-M8, Mutant MnCl_2_- 25, 50, 75, 100 µM with 10 mM MgCl_2_ ;A1-A3, Mutant dATP- 0.15, 0.20, 0.25 mM with 1.0 mM other dNTPs ; G1-G3, Mutant dGTP- 0.5, 0.75, 1.0 mM with 0.1 mM other dNTPs ; MG1-MG2, Mutant with 5, 10 mM MgCl_2_ ;D1-D2, Mutant with DMSO (V/V) conc. 5, 10 %)

| **Sample** | **Crude enzyme (ml)** | **Casein (0.6%w/v) (ml)** | **Buffer volume (ml)** | **Absorbance**  **(280 nm)** | **Crude enzyme activity (IU)** |
| --- | --- | --- | --- | --- | --- |
| B.B | 0 | 0 | 3.5 | 0.001 | - |
| E.B | 0 | 3 | 0.5 | Auto zeroed | - |
| S.B | 0.5 | 0 | 3 | 0.002 | - |
| **WT** | **0.5** | **3** | **0** | **0.632** | **14.85844** |
| M1 | 0.5 | 3 | 0 | 0.632 | 14.85844 |
| M2 | 0.5 | 3 | 0 | 0.634 | 14.90995 |
| M3 | 0.5 | 3 | 0 | 0.631 | 14.83268 |
| M4 | 0.5 | 3 | 0 | 0.632 | 14.85844 |
| M5 | 0.5 | 3 | 0 | 0.633 | 14.88419 |
| M6 | 0.5 | 3 | 0 | 0.631 | 14.83268 |
| M7 | 0.5 | 3 | 0 | 0.623 | 14.62664 |
| M8 | 0.5 | 3 | 0 | 0.634 | 14.90995 |
| **A1** | **0.5** | **3** | **0** | **3.010 (after 7 times dilution)(Abs.=0.623)** | **114.151** |
| A2 | 0.5 | 3 | 0 | 0.632 | 14.85844 |
| A3 | 0.5 | 3 | 0 | 0.641 | 15.09024 |
| G1 | 0.5 | 3 | 0 | 0.002 | -1.36763 |
| G2 | 0.5 | 3 | 0 | 0.621 | 14.57512 |
| G3 | 0.5 | 3 | 0 | 0.618 | 14.49786 |
| D1 | 0.5 | 3 | 0 | 0.651 | 15.34779 |
| D2 | 0.5 | 3 | 0 | 0.631 | 14.83268 |
| MG1 | 0.5 | 3 | 0 | 0.721 | 17.15069 |
| MG2 | 0.5 | 3 | 0 | 0.654 | 15.42506 |

**Table S3a	Screening of the *rsep* mutant with enhanced protease activity**

**Table S3b	Comparison of the relative protease activities of the *rsep* mutants with *rsep* recombinant wild-type**

| **Mutants** | **Relative protease activity** |
| --- | --- |
| WILD TYPE | SAME |
| M1 | COMPARABLE TO WT |
| M2 | COMPARABLE TO WT |
| M3 | COMPARABLE TO WT |
| M4 | COMPARABLE TO WT |
| M5 | COMPARABLE TO WT |
| M6 | COMPARABLE TO WT |
| M7 | COMPARABLE TO WT |
| **A1** | **7.70 TIMES INCREMENT*** |
| A2 | COMPARABLE TO WT |
| A3 | COMPARABLE TO WT |
| G1 | LOST***** |
| G2 | COMPARABLE TO WT |
| G3 | COMPARABLE TO WT |
| MG1 | SLIGHT INCREASE |
| MG2 | SLIGHT INCREASE |
| D1 | SLIGHT INCREASE |
| D2 | COMPARABLE TO WT |

**Table S4:	Specific activity of *rsep* metalloprotease mutant A1**

| **Substrate** | **Wild-type metalloprotease**  **(IU/mg)** | **Mutant metalloprotease**  **(IU/mg)** | **Improvement in protease activity of mutant** |
| --- | --- | --- | --- |
| **Casein** | 6.31 | 50.68 | 8.04 times |
| **Azocasein** | 6.30 | 49.90 | 7.92 times |
| **Gelatin** | 5.94 | 46.63 | 7.85 times |

**Table S5	Comparison of the characteristics of the *rsep* mutants with *rsep* recombinant wild-type**

| **Characteristics** | ***rsep w*ild-type (WT) metalloprotease** | ***rsep*A1 *m*utant metalloprotease** |
| --- | --- | --- |
| **pH optima** | 9.0 | 8.5 |
| **Temp. optima** | 35 °C | 40 °C |
| **pH stability** | Stable at alkaline pH (7.5-10.0) for 48 h | Stable at alkaline pH (8.0-10.5) for 48 h |
| **Temp. stability** | Comparatively less stable at higher temperatures | Comparatively more stable at higher temperatures |
| **Molecular weight** | 46 kDa | 92 kDa |
| **Stable Native form** | Monomer | Dimer |
| **Specific Activity** | 7.24 IU/mg | 50.68 IU/mg |
| **Substrate specificity** | Less broad specificity | More broad specificity |
| **Catalytic properties** | Km =391.53 µM; Vmax.=29.59 µM/mL/min | Km =185.867 µM; Vmax.=30.74 µM/mL/min |
| **Melting temperature** | 62.73 °C | 70.43 °C |
| **Relative catalytic efficiency** | The relative catalytic efficiency of mutant is 4.208 times as compared to the wild type | |
| **Stability towards inhibitors** | Stable in almost all inhibitors except for EDTA, 1,10 Phenanthroline | Stable in almost all inhibitors except for EDTA, 1,10 Phenanthroline |
| **Stability towards metal ions** | The protease was stable in almost all the metal ions except for Hg.  Zn^2+^ is essential for activity | The protease was stable in almost all the metal ions except for Hg.  Zn^2+^ is essential for activity |
| **Stability towards detergents** | Stable in almost all detergents except for Beta Mercaptoethanol, and DTT. It was feebly active in SDS and Urea | stable in almost all detergents except for Beta Mercaptoethanol, and DTT. It was feebly active in SDS and Urea |
| **Stability towards organic solvents** | Stable in tetradecane, dodecane and decane upto 24 h  Stable in heptane and octane for 6 h; when incubated for 12 h, residual activity decreases to 82%, while in 24 h, the residual activity decreases to 66%.  Isooctane, hexane, cyclohexane, and benzene followed similar trend.  The protease remain unstable in Toluene, DCM, and Butanol | Completely stable in tetradecane, dodecane, and decane up to 24 h. In heptane and octane, the proteases are stable for 6 h, but when incubated for 12 h, the activity decreases to nearly 80% which again decreases to 65% when incubated for 24 h. A similar trend is followed when incubated in isooctane, hexane, cyclohexane, and benzene for a total of 24 h. The stability of the proteases perishes in the presence of toluene, DCM, and butanol. |
| **Tertiary structure**  **(Fluorescence spectra)** | The comparative fluorescence spectra showed decreased intrinsic fluorescence intensity of the *rsep* mutant metalloprotease compared to the wild-type *rsep* metalloprotease. This inferred the appearance of new polar residues or replacement of amino acid residues or both in the primary structure of the *rsep* mutant metalloprotease. | |
| **Secondary Structure**  **(CD spectra)** | The CD spectra concluded the decrement in the percentage of α-helices compared to *rsep* WT, which indicated substitution of alanine residues, which has the highest propensity of forming alpha helices, in the primary structure of *rsep* mutant metalloprotease. | |
| **Nucleotide sequence analysis** | This showed the appearance of new polar amino acid residues in the primary sequence of the *rsep* mutant metalloprotease compared to the *rsep* WT metalloprotease. These amino acids were chiefly Histidine, Lysine, Glutamic acid, and Aspartic acid. The data presented the highest increase in the number of Histidine residues | |
| **ICP-MS analysis** | Results indicated 3.03 times more uptake of zinc ions in case of mutant compared to the wild type-*rsep* which correlates with the results of sequencing, effects of Zn^2+^ metal ions, and fluorescence data that described the increment in the histidine residues that interact with zinc ions in such metalloproteases which resulted in enhanced protease activity and affinity of the evolved mutant *rsep* metalloprotease through directed evolution | |
